# Computationally Designed RNA Aptamers Enable Selective Detection of FUS Pathology in ALS

**DOI:** 10.1101/2025.04.30.651570

**Authors:** Elsa Zacco, Martina Gilodi, Abigail Dos Santos, Alexandros Armaos, Rachel James, Francesco Di Palma, Fergal M. Waldron, Marco Scotto, Jakob Rupert, Vlad Korobeynikov, Neil A. Shneider, Mathew H. Horrocks, Jenna M. Gregory, Gian Gaetano Tartaglia

## Abstract

FUS is an RNA/DNA-binding protein whose mislocalization and aggregation are defining pathological features of amyotrophic lateral sclerosis (ALS) and frontotemporal lobar degeneration (FTLD). Detecting pathological FUS assemblies remains challenging, as antibody-based approaches are frequently limited by epitope masking, conformational heterogeneity, and cross-reactivity with physiological FUS pools.

Here we apply rationally designed RNA aptamers to selectively recognize FUS across soluble and aggregated states. The aptamers bind solvent-exposed RNA-binding regions of FUS with low-nanomolar affinity and adopt stable hairpin conformations that support high specificity. In cultured cells expressing the ALS-associated p.P525L FUS mutation, the aptamers detect cytoplasmic and nuclear FUS assemblies that are frequently missed by commercial antibodies and show reduced recognition of FUS-containing protein complexes.

Using super-resolution imaging, the aptamers enable visualization of early FUS aggregation intermediates that are inaccessible to conventional amyloid dyes. In post-mortem brain tissue from individuals with FUS-ALS, aptamer staining selectively labels pathological FUS while sparing normal nuclear FUS, revealing prominent nuclear and nucleolar pathology that is poorly resolved by antibody-based methods.

Together, these findings establish RNA aptamers as sensitive and selective probes for pathological FUS and uncover previously underappreciated features of FUS aggregation in ALS. This work highlights the value of nucleic-acid–based recognition tools for interrogating protein misfolding and neurodegenerative disease pathology.

## INTRODUCTION

FUsed in Sarcoma (FUS), also known as Translocated in LipoSarcoma (TLS), is a multifunctional DNA/RNA-binding protein with significant roles in maintaining cellular homeostasis through its involvement in RNA metabolism, stress response, gene expression, and DNA repair ^1^. FUS shuttles between the nucleus and the cytoplasm, and this dynamic localization is crucial for its function in both RNA metabolism and response to cellular stress ^2^. Through its interaction with RNA, FUS regulates splicing, transport, stability, and translation and it is involved in the processing of pre-mRNA into mature mRNA ^3^.

Mutations or dysregulation of FUS protein have been linked to several neurological disorders, particularly neurodegenerative diseases. For example, mutations in the FUS gene have been identified in a subset of familial amyotrophic lateral sclerosis (ALS) cases ^4^ and alterations in the FUS gene have also been associated with certain forms of frontotemporal lobar degeneration (FTLD) ^5,6^. In these and other pathologies, the physiological functions of FUS are compromised by the accumulation of the protein in toxic aggregates, insoluble structures in which FUS transit from a liquid state to a solid-like form ^7^. FUS aggregation –or liquid-to-solid phase transition (LSPT)– is thought to contribute to disease pathogenesis particularly with regards to ALS and FTLD, for which the relocation of FUS from the nucleus to the cytosol and the consequent cytoplasmic protein accumulation are considered pathological markers of disease ^8,9^. The exact molecular mechanism underlying FUS aggregation and the way FUS LSPT correlates to disease development is still unknown. Various factors –including genetic mutations, changes in cellular stress response, impaired protein quality control mechanisms, and disruption of RNA metabolism– are believed to contribute to FUS aggregation and its pathological consequences ^10,11^ .

Efforts to accurately detect and track FUS aggregation are increasingly critical for timely diagnosis and therapeutic intervention in ALS–FTD. Current antibody-based approaches for visualising FUS pathology are inherently limited. First, commercially available antibodies raised against FUS exhibit variable cross-reactivity with other members of the FET protein family, including EWSR1 and TAF15, complicating the attribution of detected signal specifically to FUS ^12^. Second, different FUS antibodies result in markedly staining patterns with distinct localisation patterns, ranging from predominantly nuclear staining to cytoplasmic aggregates or diffuse signal, with further discrepancies observed between biochemical assays and imaging-based modalities ^13^. These inconsistencies likely reflect both differences in epitope recognition and the structural heterogeneity of FUS assemblies. Third, the accurate detection of FUS pathology has never been more therapeutically relevant, as emerging FUS-targeted interventions advance toward clinical translation ^14^. Finally, antibody-based methods may fail to capture clinically meaningful FUS pathology altogether. Our recent work demonstrated that FUS is present within pathological aggregates in sporadic ALS and SOD1-associated cases, where it co-aggregates with TDP-43, pathology that may be overlooked using conventional immunodetection strategies ^15^. Together, these findings underscore the urgent need for improved tools to detect disease-relevant FUS species with greater specificity and consistency, beyond what is currently achievable with available antibodies.

There is therefore a need for an alternative means by which to target and track FUS aggregation with higher accuracy. In this context, employing RNA-based aptamers to interact and engage with FUS aggregates might represent a promising solution. Aptamers are short single-stranded oligonucleotides that fold into a particular three-dimensional structure and that bind a target molecule with high specificity ^16,17^ . Using aptamers, a combined effect of structural and sequence recognition could guarantee an affinity towards the target within the picomolar-nanomolar range. The small size —usually between 5 and 15 kDa— and their chemical features give several advantages to the aptamers over antibodies in terms of tissue penetration ability, chemical stability and solubility properties. Their nature (DNA or RNA) renders them non-immunogenic, chemically versatile and synthetically accessible.

In the case of FUS, attempts have been made to narrow down the RNA sequences and structures with which this protein might show preferential interaction. The literature presents a few cases in which, taking advantage of its natural ability to bind to nucleic acids, some oligonucleotide sequences have been identified to interact with FUS with certain specificity and have been employed mostly to clear mechanistic details of FUS binding to its RNA partners. Sequential Evolution of Ligands by Exponential Enrichment approach (SELEX) ^18^ revealed that FUS displays stronger binding avidity towards RNA sequences containing the GGU motif ^19^, while studies on the structural preferences of FUS showed that the protein interacts favorably with hairpin RNAs or with G-rich sequences forming the G-quadruplex quaternary structure ^20,21^. Moreover, nuclear magnetic resonance (NMR) experiments demonstrated that both RRM and Zn finger domains of FUS take concomitant part to the binding of RNA. While the first motif is most likely to be involved in the binding with the structural components (hairpins/G-quadruplexes), the latter confer the protein its preference towards GGU-rich RNA sequences ^22^.

With this knowledge at hand, the possibility of tailor-designing RNA sequences to selectively bind FUS appears more accessible, although it remains challenging to achieve using currently available experimental approaches. In this work, we offer a more approachable in silico strategy for RNA aptamer selection against FUS and demonstrate the utility of the resulting aptamers in tracking FUS aggregation in vitro, in cellular systems, and ex vivo. By enabling sensitive and selective detection of pathological FUS across multiple biological contexts, this approach paves the way for the development of new diagnostic tools for FUS-driven neurodegenerative diseases.

Beyond diagnostics, the generation of RNA aptamers that selectively bind FUS opens multiple downstream applications, including: (i) targeted therapeutic strategies, such as blocking pathogenic FUS–RNA or FUS–protein interactions or delivering therapeutic cargos specifically to cells containing pathological FUS; (ii) biomarker detection platforms, through incorporation of FUS-binding aptamers into imaging tracers (e.g. PET or MRI) or affinity-based assays for sensitive and specific detection of FUS levels, subcellular localization, or aggregation state in patient samples; and (iii) the development of novel research tools, such as aptamer-based pull-down reagents, live-cell imaging probes, or modulators of FUS phase behavior, enabling mechanistic studies of FUS function and dysfunction in RNA metabolism and neurodegeneration.

## RESULTS

### A computational pipeline for the *in silico* design of RNA aptamers targeting FUS

One of the main goals of this work was to extend a computational aptamer design strategy previously introduced and validated for other proteins ^17,23,24^, adapting it to the specific challenges posed by FUS aggregation. Building on catRAPID-based interaction modeling ^25,26^, we modified the design pipeline to account for domain accessibility and aggregation-associated structural constraints, thereby leveraging the intrinsic nucleic acid-binding properties of FUS. This refined in silico framework enabled efficient prioritization of RNA sequences with high predicted affinity and specificity, substantially accelerating the identification of aptamers suited for targeting pathological FUS assemblies.

Specifically, aptamers were designed to interact with the RNA-binding portion of FUS, encompassing amino acids 269–454 (hereafter referred to as FUS_269–454_). FUS_269–454_ includes the RRM, Zn finger, and one of the three RGG domains, all elements implicated in RNA interactions ^22,27^. The aptamer candidates were selected based on three principal criteria: (i) the presence of enriched sequence motifs derived from FUS CLIP datasets; (ii) the ability to adopt a stable hairpin fold, favoring binding to the RRM domain ^20^; and (iii) the presence of a GGU triplet downstream of the stem-loop, which facilitates interaction with the Zn finger domain of FUS ^22^. Most importantly, we have previously shown that protein aggregation sequesters the amyloid core while leaving hydrophilic, RNA-interacting regions solvent-exposed, providing accessible surfaces that are amenable to targeting by RNA aptamers ^26^. Consistent with this model, aggregation propensity analysis of FUS shows that the RNA-binding regions included in FUS_269–454_ are predicted to be external to the aggregation core and thus available for aptamer binding even in aggregated states (Suppl. Fig. 1). This key property underlies the success of targeting FUS aggregates with RNA aptamers. Furthermore, alphaFold-based modeling of multimeric FUS assemblies predicts an aggregated N-terminal core and solvent-exposed RNA-binding regions, including the RRM and RGG2, consistent with their accessibility for aptamer binding (Suppl. Fig. 1; Materials and Methods).

From ENCODE eCLIP data (HepG2 and K562), 4,203 high-confidence FUS-binding sites were identified (fold enrichment > 3). Of these, 401 overlapped with PAR-CLIP data and were used as the positive set for motif discovery, while 1,075 control sequences were sampled from low-confidence, non-overlapping eCLIP peaks (Fig. 1A). catRAPID effectively distinguished FUS-binding from non-binding RNA regions identified by eCLIP (AUC = 0.868; Fig. 1B), supporting its suitability for prioritizing RNA sequences with high interaction potential for FUS ^25^. Motif analysis identified 11 FUS-recognized sequence motifs, which were used to generate 28,102 candidate aptamers with 0–20 nt 31 extensions (Materials and Methods).

**Figure 1:**
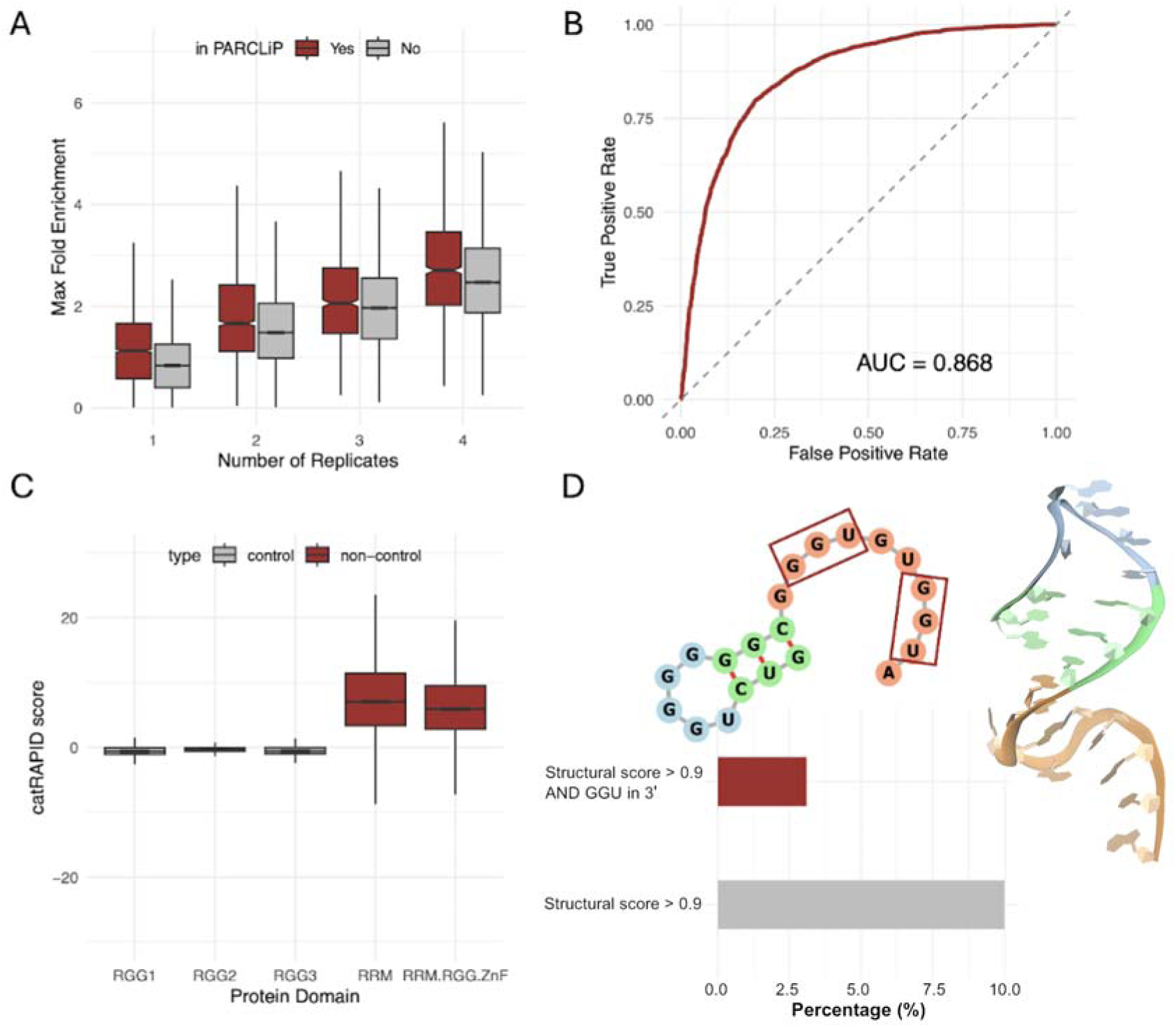
Conceptual framework for rational RNA aptamer design targeting FUS. **A:** Boxplot showing the distribution of maximum fold enrichment for binding sites detected across one to four biological replicates. Sites overlapping with PAR-CLIP data (red) show higher fold enrichment compared to those not overlapping (grey); **B:** Receiver operating characteristic (ROC) curve evaluating the ability of catRAPID to discriminate true FUS-binding sites from controls. The area under the curve (AUC) is 0.868, indicating strong predictive performance; **C:** Boxplot illustrating the catRAPID interaction scores between candidate aptamers and different FUS domains. Sequences targeting the RRM and RRM.RGG.ZnF domains (red) show significantly higher predicted interaction propensity compared to controls (grey), whereas RGG-only domains show low interaction scores; **D:** Summary of final aptamer selection. Among the sequences with high secondary structure scores (>0.9), a subset containing the GGU motif in the 3’ extension (red) was prioritized for experimental validation. An illustrative example of both the predicted secondary and tertiary structures of a selected aptamer is shown, highlighting the prototypic hairpin conformation and the location of GGU motifs.

We then assessed domain-specific binding propensities by computing catRAPID scores across our full pool of 28,102 candidate sequences. Median interaction scores for the RGG1, RGG2, and RGG3 domains were –0.66, –0.36, and –0.60, respectively, whereas the RRM and the combined RRM + RGG + ZnF region scored 7.04 and 5.89 (Fig. 1C), confirming a strong predicted affinity of our motifs for the canonical RNA-binding surfaces.

A scoring function was developed to evaluate secondary structures ^28^, prioritizing those with a perfect hairpin loop at the 5’ end, followed by a linear, unstructured sequence at the 3’ end ^22^. A total of 1,463 aptamers achieved an optimal secondary structure score (SS_score = 1), and 2,613 aptamers scored above 0.9 (SS_score > 0.9). Among these, 605 aptamers contained at least one GGU subsequence in the 3’ extension and none GGU subsequence in the hairpin loop. These aptamers were finally ranked for their predicted interaction score with FUS_269-454_ defining a chart of most promising candidates; the predicted secondary and tertiary structures of one of the selected aptamers resulting in that prototypic hairpin-loop structure with a 3’ extension containing the GGU motif are shown in Fig. 1D. A list of negative controls was also compiled by ranking CLIP data regions proximal to the interaction peaks according to their negative interaction score. Among these, we selected the ones not containing the triplet GGU and with a hairpin score = 0. The top ranked negative aptamer (worse interaction score) was selected for these studies as a control (Materials and Methods).

### Experimentally determined binding strengths agree with computational predictions

To verify the accuracy of the computational pipeline to identify RNA aptamer candidates targeting FUS with good interaction strength, a series of RNA sequences have been selected to experimentally determine their binding strength towards FUS_269-454_. The dissociation constant (K_d_) was used as a measure of interaction strength between the RNA and the protein and was determined via biolayer interferometry (BLI), an optical technique to measure interactions between molecules. From the aptamer list generated computationally, both the most promising candidates and a few controls derived from intermediate steps of the aptamers development were tested. The aptamers were screened experimentally. The six aptamers derived from the latest steps of the computational design pipeline are indeed the ones displaying the strongest affinity towards FUS_269-454_, with a K_d_ ranging from 23 nM to 740 nM (Fig. 2A-C and Suppl. Fig. 2A-D). The two aptamers deriving from intermediate steps of the pipeline, extracted at a stage in which they were not yet scored for their hairpin structure, display a slightly worse K_d_ of 1100 and 1900 nM (Suppl. Fig. 2E-F).

**Figure 2:**
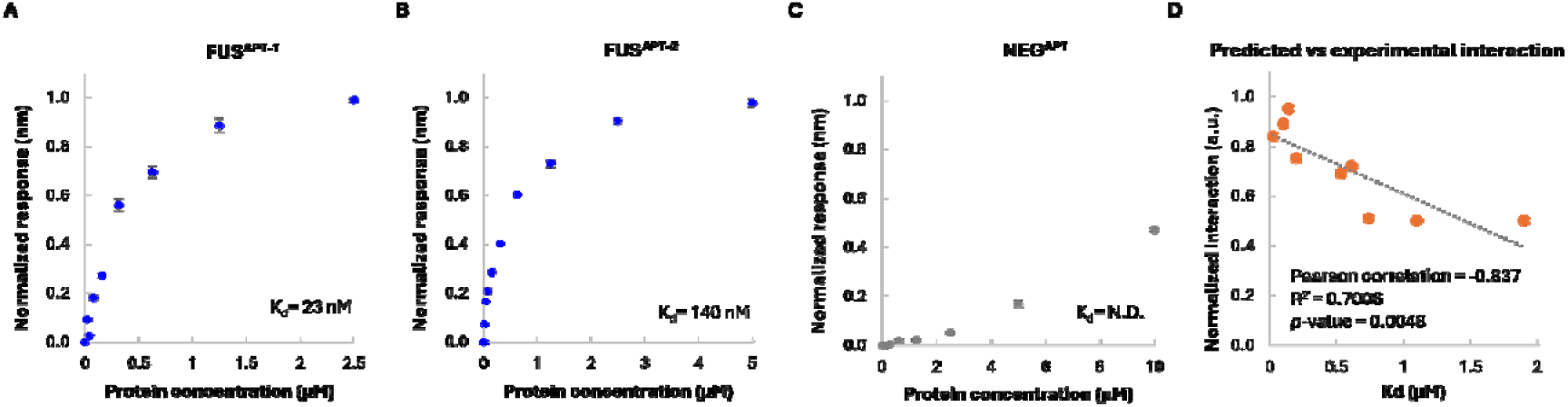
Binding of selected aptamers with FUS_269-454_. **A-C:** Biolayer interferometry-generated binding curves, showing the normalized response as a function of the protein concentration, from which the dissociation constants (K_d_s) were determined. Binding curves for FUS^APT-1^, FUS^APT-2^ and for the negative control NEG^APT^ are reported (N.D.= not determined). **D:** Comparison between predicted interaction scores and experimentally determined K_d_s of the nine aptamers with the highest scores, showing a significant negative correlation (Pearson correlation= −0.837).

Taken together, there is an excellent correlation between the predicted interaction scores and the experimentally measured K_d_ (Pearson correlation = −0.837, Fig. 2D).

To further verify the superiority of the in silico-designed best aptamer candidates compared to other RNA sequences developed by others using different means, the K_d_ between FUS_269-454_ and other aptamers found in literature was determined. Two aptamers, here called ComboNMR17 and ComboNMR22, were derived from RNA sequences deposited in the protein data bank (PDB) in association with either the RRM ^29^ or the ZnF of FUS ^30^. The RNA oligonucleotides were combined together to generate aptamers that could potentially bind both domains of FUS. In both these cases, a K_d_ of 1600 nM was determined (Suppl. Fig. 2G-H), making these aptamers comparable to the ones generated during the intermediate steps of the computational pipeline.

The top aptamer candidates were also compared to other three RNA oligonucleotides reported in literature to bind FUS (Suppl. Fig. 2I-K). The RNA named PrD was chosen because, as the aptamers designed in silico, it folds into a stable hairpin and it contains the GGU motif on its tail ^31^. The other two aptamers, named PSD95GQ2 and Shank1A, instead share the ability to fold into G-quadruplexes^21^. This structural feature was not preferred during the development of the in silico-disigned sequences because of its known promiscuity of binding ^32^. Nevertheless, the affinity of these sequences towards FUS was reported to be quite high. For these three RNA aptamers, the K_d_ reported in the literature was determined by methods different from BLI and the affinity was tested towards a different FUS-derived construct compared to FUS_269-454_. For these reasons, the binding strength of these sequences was again determined using the same conditions as the ones employed to calculate it for the in silico-generated aptamers (e.i. using BLI to derive the K_d_ with FUS_269-454_).

The K_d_ determined by BLI for PrD was 2100 nM (Suppl. Fig. 2I), placing this aptamer in the same range of the controls ComboNMR17 and ComboNMR22, with which PrD shares the hairpin structure. PSD95GQ2 and Shank1A displayed instead a K_d_ with FUS_269-454_ of 220 nM and 713 nM, respectively (Suppl. Fig. 2J-K). These binding strengths are comparable to some of the RNA aptamers generated via the computational pipeline but are not as strong binders as the top four RNA sequences generated in silico.

To conclude the interrogation of possible RNA aptamer controls, a sequence of length comparable to the most promising candidates, lacking both the hairpin structure and the GGU motif, was selected (for their design, see Material and Methods section). For this sequence, named NEG^APT^, it was not possible to determine a K_d_ for FUS_269-454_, since no binding occurred in the selected conditions (Fig. 2C).

In conclusion, the RNA aptamers designed through the computational pipeline described above are superior or equal to other RNA sequences reported in the literature in binding the RNA binding region of FUS comprising the RRM, the ZnF and one RGG repeat. Having been generated in silico, these RNA aptamers present the additional advantage of a faster and cost effective production compared to others. The use of computational tools also enables the screening of sequences that not only are predicted to strongly bind FUS but also to target it with high specificity, by selecting RNA aptamers predicted to prefer FUS to other proteins of similar composition and length.

### Circular dichroism and structural predictions confirm the presence of stable hairpins in designed aptamers

One of the main features of the aptamers designed with the computational pipeline was a high hairpin score. For this reason, the two most promising candidates, FUS^APT-1^ and FUS^APT-2^, were investigated for their ability to fold into this structure by circular dichroism (Fig. 3A-C). Together with the two best performing aptamers showing the strongest binding strength towards FUS_269-454_ (Fig. 3A-B), also the negative control NEG^APT^ was analyzed (Fig. 3C). Alongside the experimental determination of the aptamers secondary structures, their RNAComposer ^33,34^ structure predictions are also reported (Fig. 3D-F).

**Figure 3:**
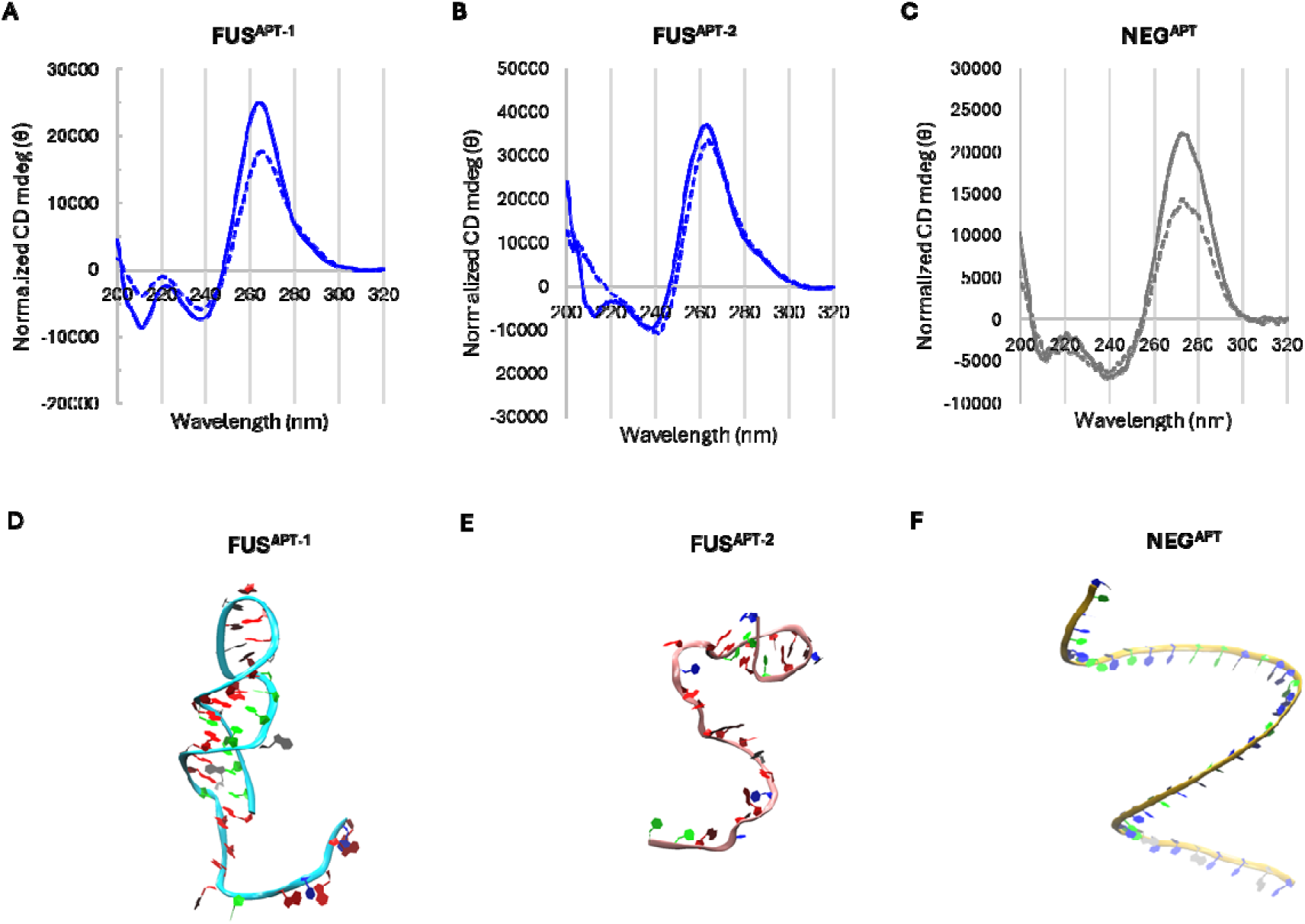
Structural analysis of selected aptamers. **A-C:** Circular dichroism profiles of FUS^APT-1^, FUS^APT-2^ and NEG^APT^, highlighting the structural stability and complexity of the first two compared to the negative control. **B:** RNAComposer predictions of the same aptamers, showing the three-dimensional structures modelled for FUS^APT-^ and FUS^APT-2^, and the fully single stranded nature of NEG^APT^.

FUS^APT-1^ and FUS^APT-2^ share a similarly-shaped CD profile, with a main maximum at 265 nm and a minimum at 240 nm (Fig. 3A-B). These features are generally indicative of an RNA sequence, and indeed we find these characteristics also in the CD profile of NEG^APT^, albeit in this case the maximum is shifted towards higher UV regions (273 nm) (Fig. 3C). It is also observable a difference in the intensities of the peaks between FUS and FUS : position and amplitude of the peaks may markedly differ with sequence, not only because the chromophore differ but also because of the different conformational properties of each base. What is important to observe for this study is the presence of a weak shoulder at 285-290 nm, which is a typical feature indicative of a hairpin . This shoulder is absent in the CD profile of NEG^APT^.

To confirm the stability of the hairpins in the selected conditions, the CD spectra were acquired at increasing temperature and the CD profiles at 30 °C were compared with the ones recorded at 70 °C (Fig. 3). For the same sequence at the same concentration, a loss of intensity is indicative of an increased instability of the RNA. FUS and FUS display a loss of intensity of the maximum at 265 nm of around 1.2 folds, while the shoulder indicative of the hairpin remains unvaried in all cases (Fig. 3A-B). Instead, the maximum at 270 nm of NEG^APT^ drops 1.6 times from its initial intensity (Fig. 3C). This suggests strong secondary structure features for the two top aptamers and the lack of any stable configuration for NEG^APT^.

Querying the RNAComposer server with the FUS^APT-1^, FUS^APT-2^ and NEG^APT^ corresponding sequences, we obtained their 3-dimensional models (Fig. 3D-F). The predicted tertiary structure for FUS is a 7-base-pair stem-loop with a 12 nucleotide-long single-stranded tail ending with the UGGUG sequence (Fig. 3D). Such a composed structure is based on different parts of several pdb structures: 4G9Z (100% homology), 1J9H (100% homology), 1VQO (100% homology), 4HUB (100% homology), 1U9S (63.6% homology), and a single strand based on 3T1Y with low homology (15.4%). Also FUS is bearing a similar structural motif in its sequence, but compared to FUS it presented a short initial hairpin (four nucleotide pairs) followed by a 16 nucleotide-long single-stranded tail (Fig. 3E); in this case, the prediction was built on parts of the following pdb structures: 3U5F (100.0% homology), 2J02 (71.4% homology) and 2XD0 (23.5% homology) for the single-strand region.

At variance with FUS^APT-1^ and FUS^APT-2^, and in agreement with the CD data, the NEG^APT^ was predicted not to have any particular secondary structure, resulting in a single-stranded model (Fig. 3F). These results indicate that the aptamers designed computationally to fold into a hairpin structure clearly display this feature in the conditions selected for this study, while the negative control, programmed to remain linear, does not show any definite structural characteristics.

### Aptamers as a super-resolution (SR) imaging probe to image *in vitro*-generated FUS aggregates

The early detection of protein aggregates in solution is essential for elucidating the molecular mechanisms underlying protein misfolding and aggregation, processes implicated in numerous neurodegenerative disorders. Conventional aggregation assays, such as those employing amyloid-binding dyes such as thioflavin-T (ThT) or ProteoStat ^17,36^, predominantly detect mature, cross-β–rich species and are therefore limited in their ability to identify early oligomeric intermediates ^37^. To overcome this limitation, we investigated whether our aptamers could be utilized for the super-resolution imaging and detection of small, early-stage protein aggregates formed by the FUS construct, FUS_269-454_.

To this end, we incubated FUS_269-454_ under conditions favouring its aggregation (see Suppl. Fig. 3A for evidence of its aggregation) and imaged the aggregates using either the amyloid-binding dye ProteoStat (Suppl. Fig. 3) or the aptamers (Fig. 4 and Suppl. Fig. 4). We collected samples from the aggregation mixture at different time-points and imaged them using aptamer-PAINT for the aptamers ^17^ (Fig. 4A-B) and standard Single Aggregate Visualisation by Enhancement (SAVE) imaging for ProteoStat (Suppl. Fig. 3B). Both FUS^APT-1^ and FUS^APT-2^ aptamers successfully enable the visualization of aggregates at a resolution of 25.44 nm ± 5.05 nm (mean ± SD, n = 56 images analysed) (Fig. 4 and Suppl. Fig. 4), as determined using Fourier Ring Correlation (FRC) ^38^, while the negative control NEG^APT^ showed minimal binding to aggregates (Suppl. Fig. 4A). At baseline, we detected 0.15 ± 0.03 aggregates/µm^2^ with FUS^APT-1^ and 0.23 ± 0.04 aggregates/µm^2^ with FUS^APT-2^ on average (Fig. 4 and Suppl. Fig. 4). As the aggregation proceeded, there was a gradual decrease in the number of detected aggregates as they slowly became larger (Fig. 4 and Suppl. Fig. 4), consistent with the detection of small oligomeric species that form rapidly and remained dispersed in the aggregation mixture.

**Figure 4:**
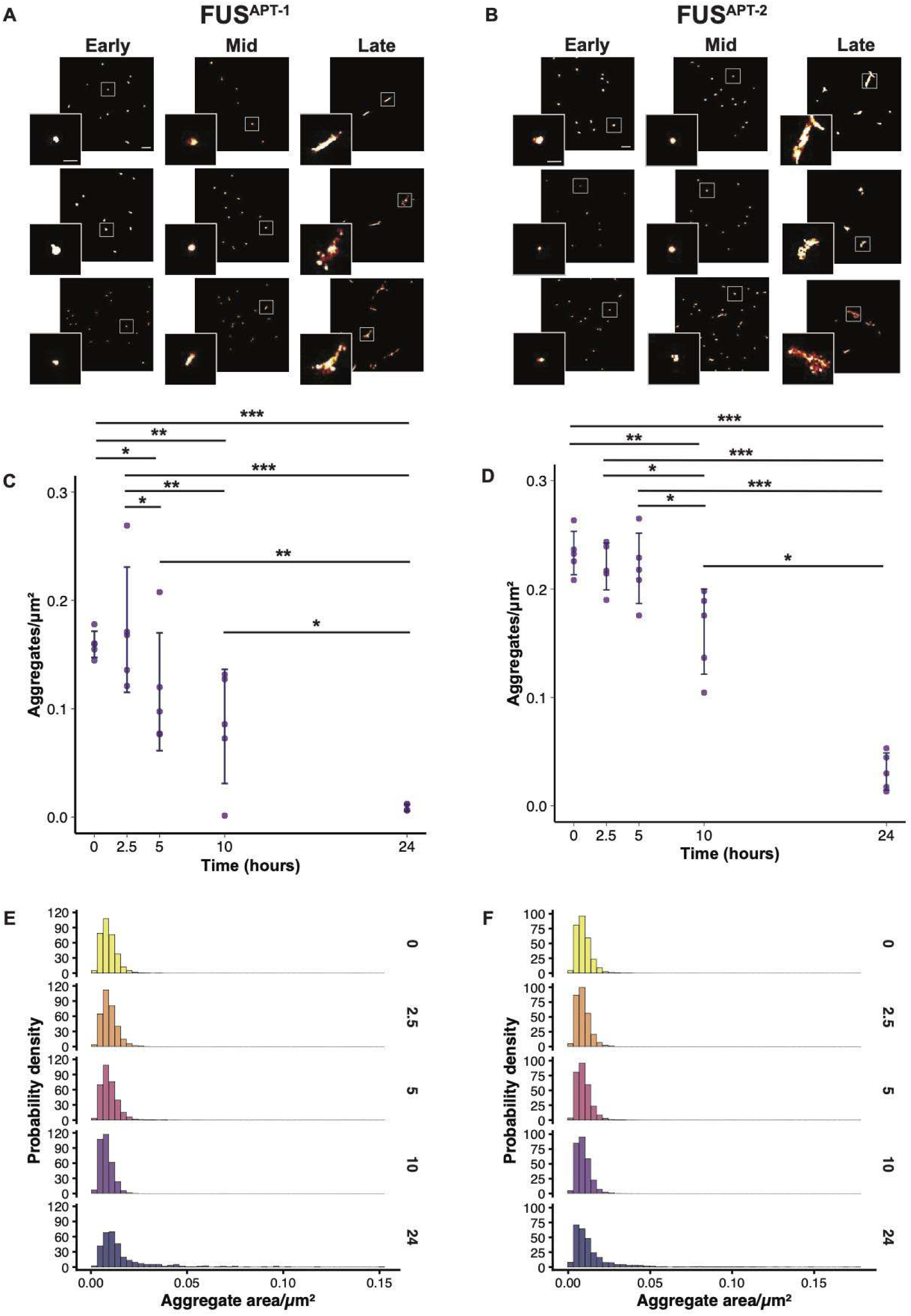
Aptamer-PAINT imaging of in vitro-generated aggregates of the FUS_269-454_ construct. Representative super-resolved images of aggregates, imaged using either **A:** FUS^APT-1^ or **B:** FUS^APT-2^. Aggregates at early stage here are from the 0 hour baseline, mid-stage aggregates are from 5 hours, and late stage from the 24 hour timepoint. Scale bars = 500 nm, scale bars on insets = 200 nm. Quantification of the numbers of aggregates detected over the aggregation period for **C:** FUS^APT-1^, or **D:** FUS^APT-2^. Each point is the average for one well of the aggregation reaction, for a total of 5 technical replicates per aptamer. Statistical evaluation (Kruskal-Wallis test, Dunn post-hoc with Benjamini-Hochberg correction) resulted in multiple significant pairwise comparisons, and adjusted p-values for these are reported in Suppl. Table 1. Normalised histograms showing the probability density distribution of areas of aggregate detected by **E:** FUS^APT-1^, or **F:** FUS^APT-2^.

At the late stage of the aggregation, there was a sharp decrease in detections with both aptamers, as the aggregates became less concentrated in the aggregation mixture and larger in size (Fig. 4 and Suppl. Fig. 4). Statistical evaluation confirmed a significant decrease in aggregates over time for both detection aptamers (Fig. 4, Suppl. Fig. 4 and Suppl. Table 1). For FUS^APT-1^, aggregate counts first begin to significantly decrease at 5 hours compared to baseline (Kruskal-Wallis test, with Dunn post-hoc, p-value = 0.035), then at 10 hours (p-value = 0.002), with an even greater reduction at 24 hours (p-value = 1.17 x 10^-7^). A similar trend was observed for FUS^APT-2^, which showed significant decreases in aggregate numbers at 10 hours (p-value = 0.008), and then at 24 hours (p-value = 3.37 x 10^-7^) compared to baseline (Fig. 4 and Suppl. Fig. 4A). To support this, there was a statistically significant increase in aggregate length at later timepoints (Fig. 4D, Suppl. Fig. 4B and Suppl. Table 1). For FUS^APT-1^, aggregate lengths were significantly higher at 10 hours after baseline (p-value = 0.001) as well as at 24 hours (p-value = 0.001). Similarly for FUS^APT-2^, significantly increased aggregate lengths were observed at 10 hours (p-value = 0.004) and 24 hours (p-value = 0.002).

Diffraction-limited imaging with ProteoStat detected consistently fewer aggregates than both aptamers, but demonstrated a similar trend (SuppI. Fig. 3). There was a significant increase from baseline to the earliest stage of aggregation assayed (Kruskal-Wallis test with Dunn post-hoc, p-value = 8.38 x 10^-6^), then a significant reduction at 10 hours (p-value = 2.61 x 10^-17^).

While Proteostat detected aggregates as early as 2.5 hours (SuppI. Fig. 3), the aptamers, which do not rely on the target having the structural motif of the β-sheet, could bind to and detect higher numbers of oligomers present in the aggregation mixture (Fig. 4 and Suppl. Fig. 4).

Together, these results demonstrate that both FUS aptamers tested enable the detection and super-resolution imaging of early, small protein aggregates that are inaccessible to conventional amyloid-binding dyes. By recognizing aggregation intermediates prior to the formation of β-sheet–rich structures, the aptamer provides a powerful tool for probing the initial stages of protein self-assembly. The ability to identify and quantify such early species holds significant potential for the development of sensitive diagnostic approaches, where detecting the onset of protein aggregation could facilitate earlier intervention in neurodegenerative disease progression.

### Aptamer-based cell staining reveals enhanced sensitivity compared to antibodies

Reliable detection of aberrant condensates and insoluble inclusions is critical for understanding and diagnosing ALS-FTD pathology. Conventional antibody-based methods can suffer from epitope masking and limited specificity, which may lead to misidentification of pathological FUS assemblies. To overcome these challenges, we tested newly developed aptamers as a sensitive and specific alternative for visualizing FUS condensates in cells.

We evaluated aptamer performance in SK-N-BE neuroblastoma cells inducibly expressing the disease-relevant P525L FUS mutant, which disrupts the nuclear localization signal, drives cytosolic mislocalization, and promotes the formation of inclusions ^39,40^. Initial Triton X-100 permeabilization produced strong nuclear aptamer signals that risked obscuring cytosolic condensates, motivating a switch to digitonin. At low concentrations, digitonin can selectively permeabilize the plasma membrane while preserving nuclear integrity ^41^. Fluorescence microscopy then was used to visualize FUS cytosolic condensates, and images were quantitatively analyzed to compare the number of condensates detected by each probe.

We compared aptamers on FUS P525L-expressing cells under oxidative stress induced by sodium arsenite, a condition commonly used to detect FUS-positive condensates ^42,43^ (Fig. 5, Suppl. Fig. 5). With digitonin permeabilization, the FUS^APT-2^ aptamer identified ∼25% more cytosolic condensates than the commercial antibody (Fig. 5). This advantage is reduced to 2% with Triton X-100 (Suppl. Fig. 6), indicating that FUS performs comparably to the antibody under harsher experimental conditions. In contrast, the FUS aptamer detected only ∼50% of the condensates labeled by the antibody (Fig. 5, Suppl. Fig. 5). Both aptamers nonetheless outperformed the negative control NEG^APT^, which stained only ∼7% of condensates (Fig. 5, Suppl. Fig. 5).

**Figure 5:**
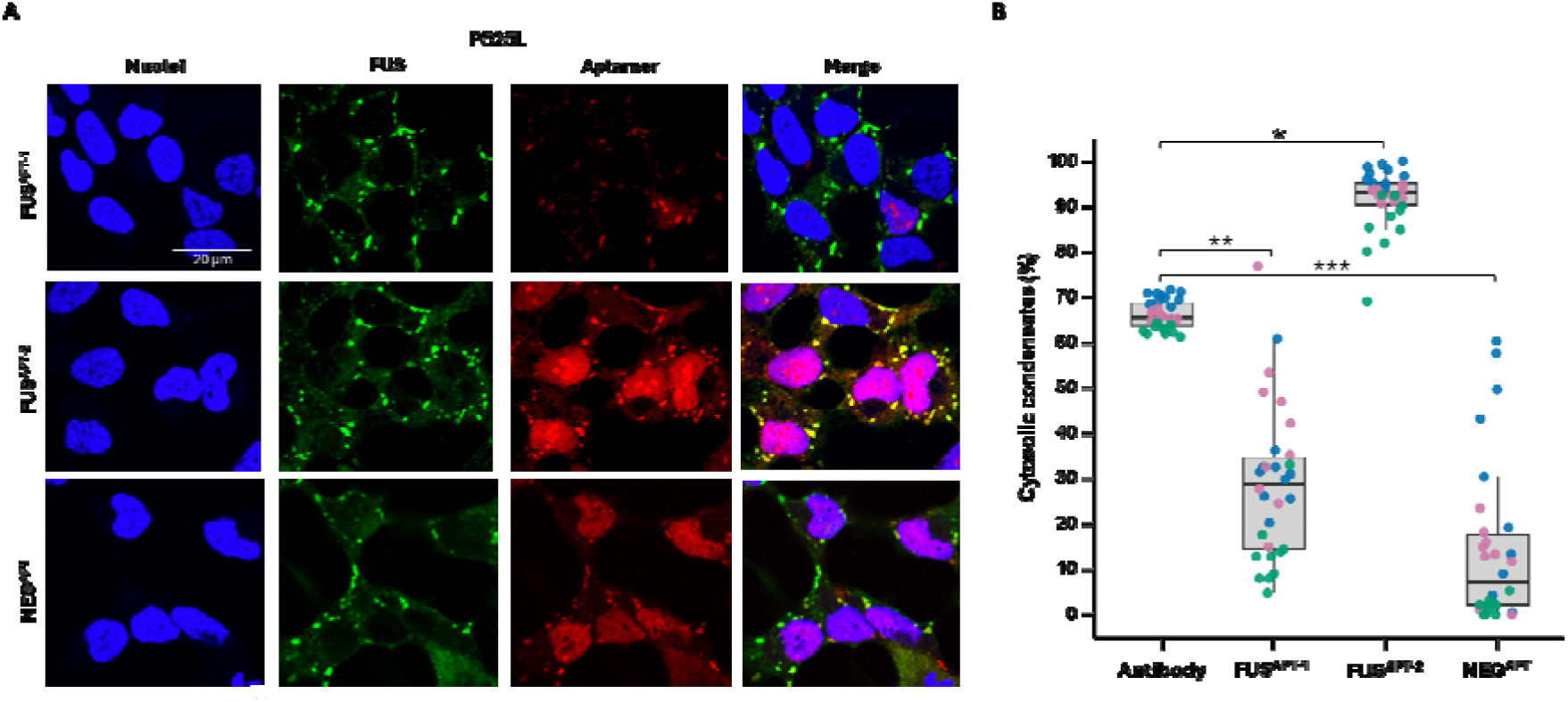
Immunostaining of FUS P525L–expressing cells subjected to sodium arsenite. Cells were stressed and stained with an anti-FUS antibody together with one of the tested aptamers. **A:** Top row: FUS^APT-1^; middle row: FUS^APT-2^; bottom row: NEG^APT^. Nuclei are labeled with DAPI (blue); the anti-FUS primary antibody is detected using an Atto488-conjugated secondary antibody (green); aptamers are labeled with an Atto590 fluorophore (red). Scale bar= 20 µm. **B:** Quantification of the percentage of cytosolic condensates detected by the antibody compared with those detected by each aptamer, normalized according to the maximum value. Points represent data from triplicate experiments (blue: replicate 1; pink: replicate 2; green: replicate 3). Statistical significance was assessed using a Nested 1-way ANOVA test with post hoc Dunnett’s multiple comparison test: * *p* = 0.0400; ** *p* = 0.0056; *** *p* = 0.000.⁵.

To complete the control panel for this experiment, we used the best performing aptamer, FUS , to analyze cells expressing wild-type FUS under basal and stress conditions alongside the mutant. In unstressed cells expressing either form of FUS, the aptamer primarily labeled the nucleus, regardless of the permeabilization method (Fig. 6, Suppl. Fig. 6), consistent with FUS known nuclear localization. A weaker, diffuse cytoplasmic signal was detected in some cells (Fig. 6, Suppl. Fig. 6), suggesting that the aptamer recognizes a largely dispersed, non-condensed FUS population in the absence of stress. Under these conditions, the antibody produced no detectable staining with digitonin (Fig. 6, Supl. Fig. 6) but labeled nuclear FUS with Triton X-100 (Suppl. Fig. 6).

**Figure. 6:**
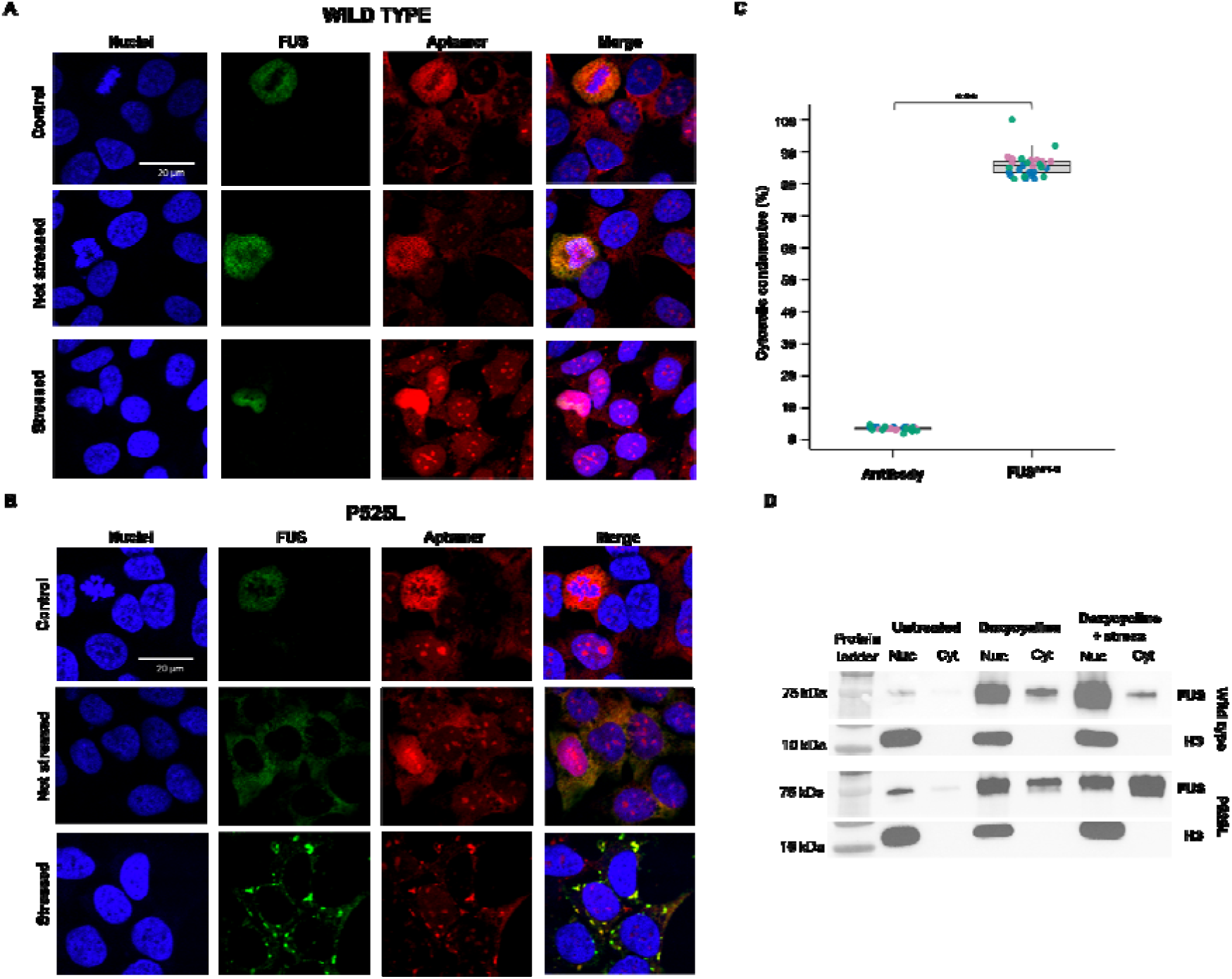
Immunofluorescence images of SK-N-BE cells permeabilized with digitonin and stained with an anti-FUS antibody together with FUS^APT-2^. A: Cells expressing wild-type FUS. **B:** Cells expressing the FUS P525L variant. For each condition, cells are shown untreated (‘Control’, top rows), following doxycycline-induced FUS expression (‘Not stressed’, middle rows), and after exposure to sodium arsenite (‘Stressed’, bottom rows). Nuclei are labeled with DAPI (blue); the anti-FUS primary antibody is detected using an Atto488-conjugated secondary antibody (green); aptamers are labeled with an Atto590 fluorophore (red). Scale bar = 20 µm. **C:** Quantification of the percentage of cytosolic condensates detected by the antibody compared with those detected by FUS^APT-2^, normalized according to the maximum value. Points represent data from triplicate experiments (blue: replicate 1; pink: replicate 2; green: replicate 3). Statistical significance was assessed using a one-tailed *t*-test: *** *p* = 0.0001. **D:** Western blots of cytosolic and nuclear fractions extracted from cells expressing either wild type (top) or FUS P525L (bottom). Histon3 (H3) is reported as housekeeping proteins for the nuclear fraction ,to confirm the absence of nuclear protein contaminations in the cytosolic fraction.

After stress induction, the antibody signal remained unchanged with either permeabilization method, showing no cytoplasmic enrichment (Fig. 6 and Suppl. Fig. 6). In contrast, the aptamer revealed distinct cytoplasmic foci in addition to the diffuse basal pattern (Fig. 6, Suppl. Fig. 6), consistent with stress-induced FUS condensates. To verify that these aptamer-positive structures contained FUS and to test whether the antibody failed to detect a conformational variant, we performed subcellular fractionation of cells expressing wild-type or P525L FUS before and after stress. Western blots showed FUS in both nuclear and cytoplasmic fractions under doxycycline-induced FUS expression, with or without stress (Fig. 6). These results indicate that the aptamer binds FUS in both compartments and stress states and can detect conformational forms inaccessible to the antibody under the tested conditions.

Together, these results demonstrate that aptamer FUS provides improved sensitivity and specificity in detecting FUS condensates compared to conventional antibodies, while FUS shows intermediate performance.

### FUS aptamers recognise aggregates in tissue more selectively than antibodies

To test the ability of selected aptamers to bind to physiologically relevant FUS aggregates in vivo, we obtained post-mortem tissue from two people who had an underlying p.P525L FUS mutation, one who had received FUS ASO therapy and one who had not^44^ and a non-neurological control. We performed immunohistochemical staining using the FUS^APT-1^ and FUS^APT-2^ sequences tagged with a 3’ biotin molecule to facilitate detection using an anti-biotin antibody immunohistochemistry method as published previously ^45^. We compared staining profiles of the two aptamers against a commercially available FUS antibody. FUS antibody staining in the non-neurological control case demonstrated a normal, predominantly nuclear staining pattern of FUS with no evidence of nuclear or cytoplasmic aggregation. In contrast to this, FUS antibody staining in the untreated p.P525L FUS mutation carrier demonstrated both normal and abnormal FUS staining including large cytoplasmic aggregates (Fig. 7A). The antibody staining in the ASO treated p.P525L FUS mutation carrier demonstrated no evidence of either normal, nor pathological FUS staining, commensurate with near complete knockdown and clearance of FUS protein. Contrastingly, the FUS aptamer staining identified only abnormal FUS staining patterns, with no staining visible in the non-neurological control and extensive pathological FUS staining in the untreated individual. Aptamer staining additionally revealed large and extensive nuclear aggregation events that are not readily identified using antibody staining alone. Similar to the antibody staining, there is no immunoreactivity to large aggregates is seen in the ASO treated individual using FUS aptamer staining, although fine cytoplasmic inclusions can be visualised by both aptamers that cannot be seen with the antibody. This staining fits with the post-mortem tissue analysis originally conducted on this material in the ASO trial, demonstrating a small amount of residual FUS in the insoluble fraction in ASO treated individuals. Finally an ASO-treated p.R521L mutation carrier was included as they were treated with ASO more than 3 half-lives (>150 days) prior to death, with the antibody staining demonstrating a reaccumulation of mostly normal FUS staining, despite the patient clinically deteriorating. The aptamers reveal that there is also a reaccumulation of large aggregates including prominent nuclear pathology. Digital image analysis was performed in QuPath using superpixel masks (Fig. 7B) ^46^. Quantification, using superpixel H-score demonstrates equivalent sensitivity of aptamer staining compared to antibody and improved specificity (no detection in control and ASO treated individuals) of aptamer staining compared to antibody. No difference was identified between the long and short FUS aptamer sequences, both demonstrated equivalent sensitivity and specificity (Fig. 7C). Amongst features readily identified by aptamer staining, nuclear pathology is most notable (Fig. 8A). Nuclear pathology is less readily visualised in antibody-stained cases as normal nuclear localisation may be obscuring the nuclear pathology. Nuclear pathological features include (i) nucleolar decoration, (ii) large nuclear inclusions, and (iii) nuclear membrane pathology (Fig. 8A). These are in addition to the large cytoplasmic aggregates, and cytoplasmic puncta that can be readily visualised by both the antibody and the aptamer sequences (Fig. 8B). No immunoreactivity is observed using the negative aptamer sequence(Fig. 8C).

**Figure 7.**
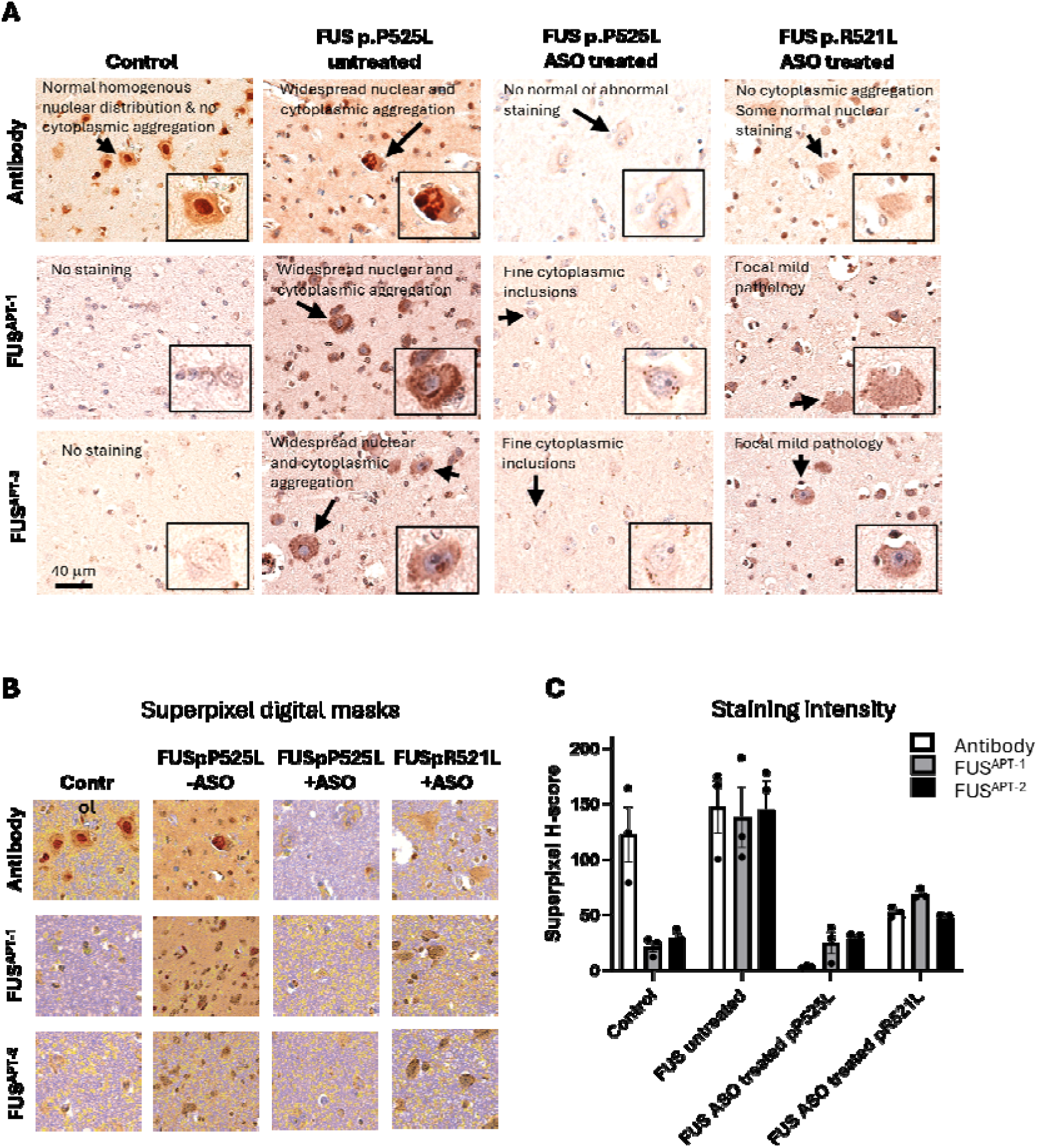
FUS aptamers recognise aggregates in tissue more selectively than antibodies. **A:** Representative photomicrographs implementing FUS antibody and aptamer staining using immunohistochemistry. Staining is performed on motor cortex from an age-matched non-neurological control, as well as from a person with an underlying p.P525L mutation who did not undergo ASO therapy, and two people with underlying p.P525L and pR521L mutations who did undergo ASO therapy. The pR521L mutation carrier received the ASO >3 half-lives prior to death and demonstrates a reaccumulation of both normal and abnormal FUS. Antibody staining demonstrates normal nuclear localisation in controls and admixed normal and abnormal FUS staining in mutation carriers including large cytoplasmic aggregates. Aptamer staining identifies only abnormal FUS staining patterns, with no staining visible in the non-neurological control and extensive pathological FUS staining in the untreated mutation-carrier. No large aggregate staining is seen in the ASO treated individual with antibody or aptamer. Aptamer staining additionally reveals large and extensive nuclear aggregation events that are not seen as frequently with the antibody staining and fine cytoplasmic inclusions in the ASO-treated cases that are not seen by antibody. **B:** Superpixel digital masks used to quantify staining intensity. Blue pixels are baseline DAB (background) and yellow, orange, and red pixels represent 1, 2 and 3 standard deviations (respectively) above baseline average intensity. **C:** Superpixel H-score used to represent staining intensity demonstrates equivalent sensitivity (compared to antibody) and improved specificity (no detection in control and ASO treated individuals). Each point is the mean of three replicates from randomly generated regions of interest within layer 5 of the motor cortex of a single patient, error bars are SD. Image analysis was performed using QuPath.

**Fig. 8.**
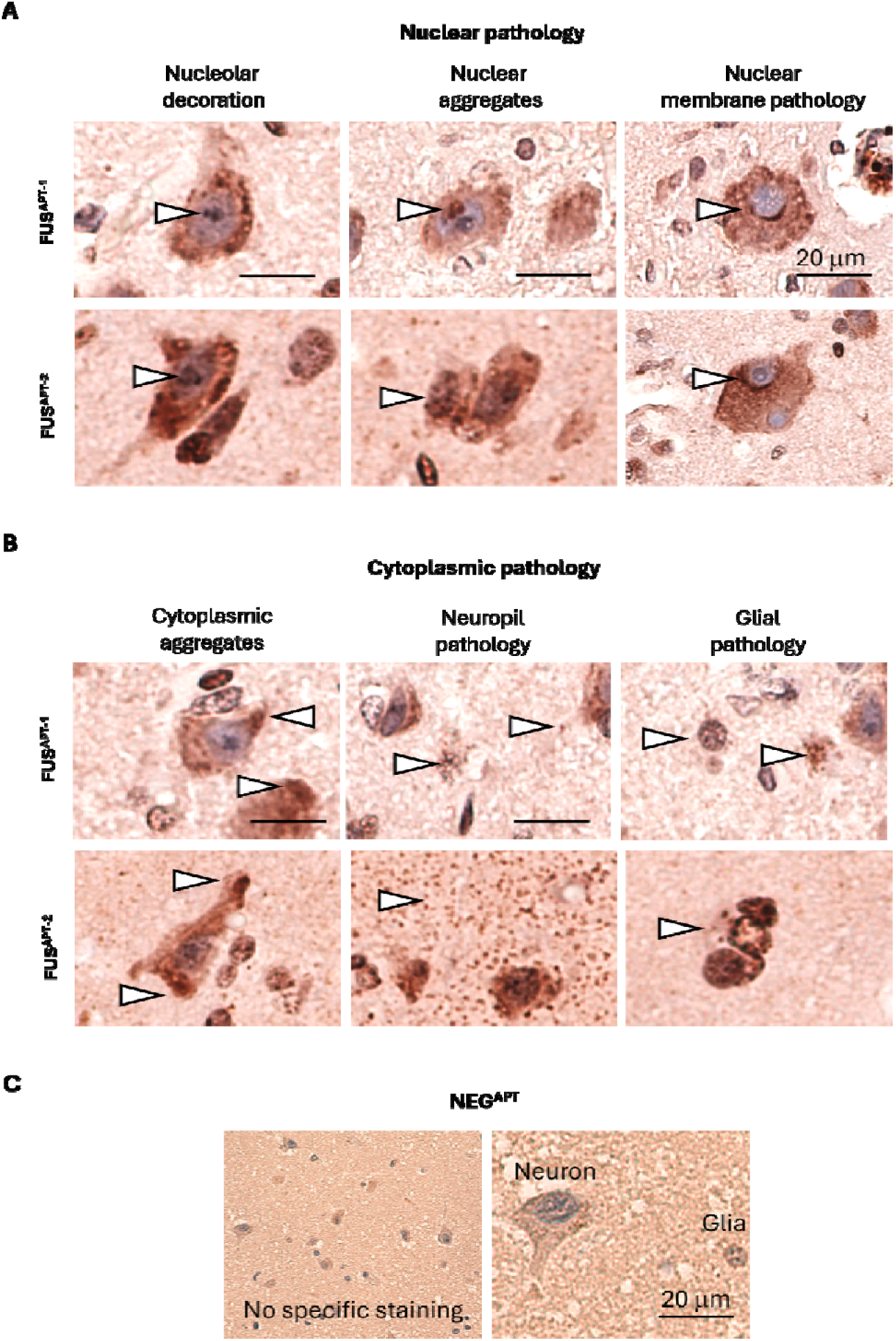
Pathological features identified using FUS aptamer histological staining. **A:** Nuclear pathology is readily visualised using aptamer immunohistochemistry. Photomicrographs demonstrate nucleolar decoration, large nuclear aggregates and nuclear membrane pathology. **B:** Cytoplasmic features show similar morphologies to antibody staining, including large cytoplasmic aggregates, neuropil staining and glial cytoplasmic puncta within processes. Features demonstrated with white arrowheads. **C:** Negative aptamer demonstrates no specific staining in neurons or glia.

## CONCLUSIONS

Aggregation of RNA-binding proteins exposes functional RNA-interacting surfaces that can be exploited for selective molecular recognition. Here, we apply a validated computational design strategy to generate RNA aptamers that engage multiple RNA-binding domains of FUS, enabling selective detection of pathological assemblies in ALS and FTLD. The need for developing RNA aptamers for FUS aggregates lies in the inherent limitations of conventional antibodies when it comes to detecting proteins that undergo state transition. Aggregated proteins often exhibit significant conformational variability—a hallmark of neurodegenerative conditions like ALS and FTLD— and antibodies typically recognize specific epitopes that may become obscured or structurally altered in aggregated states, reducing detection reliability ^47,48^. RNA aptamers possess greater structural adaptability and we can computationally tailor them for high-affinity interactions, making them well-suited to bind the diverse and often concealed surfaces of misfolded protein aggregates ^17,49^. Notably, our RNA aptamers are designed in silico, prioritizing candidates with high predicted affinity for the FUS RNA-binding region, using catRAPID interaction propensity scores alongside secondary structure predictions favoring hairpin conformations. Experimental validation confirmed that the top-ranked aptamers bind FUS with high affinity in the nanomolar range, as determined by biolayer interferometry. Circular dichroism analysis further confirmed the stability of their secondary structure under near-to-physiological conditions.

Using single-molecule localisation microscopy, we demonstrated the detection of protein oligomers and aggregates with nanometre-scale resolution at earlier stages of aggregation ^17,50,51^. At these stages, these oligomers are not accessible to conventional amyloid-binding dyes ^17,36^. Quantitative analysis revealed a progressive reduction in detectable aggregates at later time points, coinciding with a significant increase in the size of aggregates. The ability of the aptamers to detect more aggregates at earlier time points as compared to the amyloid dye ProteoStat is, in part, likely due to fundamental differences in their binding mechanisms. Amyloid-binding dyes are selective for cross-β structural motifs that arise during later stages of aggregation. In contrast, the aptamers used here engage solvent-accessible RNA-binding regions of FUS, enabling detection independently of β-sheet formation. This enables binding to structurally immature oligomers that precede fibril formation. Furthermore, at later stages of aggregation, super-resolved imaging using the aptamers were able to resolve aggregate structures more accurately than diffraction-limited imaging using ProteoStat. ProteoStat-based imaging also showed no significant difference between baseline and the 24-hour timepoint. This is likely because diffraction-limited approaches may mask the increased aggregate sizes and concomitant decrease in numbers at the late stages. In contrast, our aptamer-based approach offers increased resolution with greater precision due to their reduced signal displacement. One limitation of our approach was that aggregation was examined in vitro using the truncated FUS construct, and it is still uncertain how aptamer binding to the earliest aggregated species would be affected in the context of full-length FUS, particularly under in vitro conditions. In addition, while the two detection aptamers demonstrated consistent trends, further work will be required to establish how aptamer sequence and affinity determine sensitivity of detection during aggregation.

Cell-based experiments show that antibody detection of full-length FUS is frequently limited by epitope masking, conformational heterogeneity, and variability across fixation methods and cellular backgrounds ^13,52^. These constraints are particularly evident in stressed cells and in aggregation-prone mutants such as P525L, where nuclear and nucleolar pathology can be masked by antibody recognition of physiological FUS pools ^11,53^. In contrast, FUS^APT-2^ detects ∼25% more condensates than the antibody, including cytosolic condensates under oxidative stress that are entirely missed by immunostaining, while FUS^APT-1^ labels only a subset of antibody-positive structures. These patterns support a model in which aptamers recognize a narrower set of solvent-exposed FUS surfaces—often preserved in compact or masked conformers—whereas antibodies may bind both FUS and associated assemblies. The strong nuclear and nucleolar labeling by aptamers, particularly in P525L cells, aligns with emerging evidence that intranuclear FUS dysregulation and nucleolar involvement are early events in disease but can be overlooked with conventional immunohistochemistry ^54^. From a diagnostic perspective, aptamers offer batch consistency, structural tunability, and multiplexing compatibility, addressing several limitations of conformation-insensitive antibody workflows. At the same time, the selective performance of FUS^APT-1^ illustrates that aptamer utility remains sequence-and epitope-dependent, underscoring the need for continued optimization and benchmarking against next-generation molecular probes. The observation that FUS^APT-2^ detects cytosolic wild type FUS condensates under oxidative stress conditions that remain completely invisible to antibody staining is particularly significant. This discrepancy strongly suggests that these stress-induced condensates contain FUS conformers or packing arrangements that mask the antibody epitope while leaving the aptamer-binding surface accessible. Such a finding highlights an underappreciated conformational heterogeneity within FUS condensates and raises the possibility that widely used antibodies may systematically overlook specific, potentially transient or early pathogenic FUS species. As a result, previous assumptions about the absence or rarity of certain cytosolic FUS assemblies may need to be reconsidered. More broadly, this work underscores the value of aptamers as complementary or even superior tools for probing phase-separated or misfolded protein states, with important implications for understanding the molecular events that link oxidative stress to FUS mislocalization and the early stages of ALS/FTD-related pathology.

Importantly, post-mortem immunohistochemical analyses highlight the diagnostic potential of RNA aptamers for detecting FUS pathology, particularly in ALS cases associated with the P525L mutation, where antibody-based methods are often limiting ^40^. FUS-targeting aptamers selectively labeled pathological aggregates in untreated mutation carriers while showing no signal in non-neurological controls or ASO-treated individuals ^44^, demonstrating high target specificity in complex tissue. In contrast, antibody staining also detected physiological nuclear FUS, consistent with broader reactivity that can obscure subtle pathology. This selectivity likely reflects staining conditions that favor aptamer binding to aggregated rather than monomeric FUS, as the lower intrinsic affinity of RNA aptamers renders physiological FUS levels below the detection threshold, whereas pathological aggregation increases local epitope density and enables efficient, selective labeling of inclusions. Notably, the absence of staining in ASO-treated tissue across both aptamer- and antibody-based assays indicates that aptamer signal reflects FUS abundance rather than structural artifacts, highlighting their potential utility for monitoring therapeutic target engagement^44^. Beyond specificity, aptamer staining revealed prominent nuclear pathological features that were poorly resolved by antibodies, including nucleolar decoration, large intranuclear inclusions, and nuclear membrane abnormalities. In particular, the epitope of many anti-FUS antibodies resides at the N terminus of the protein, which is predicted to be buried within the aggregate core, particularly in the more compact nuclear inclusions. In contrast, the aptamer recognizes a broader region of FUS that is predicted to remain at least partially solvent-exposed in aggregated conformations, thereby enabling more effective detection of nuclear pathology. Quantitative analysis confirmed that aptamers match antibodies in sensitivity while offering improved specificity. Although both approaches detect cytoplasmic aggregates, the enhanced resolution of nuclear pathology positions aptamers as a powerful complement to antibody-based diagnostics, with dual-staining strategies offering a more complete assessment of FUS pathology.

In conclusion, we established a rapid, cost-effective, and scalable computational strategy for the generation of high-affinity RNA aptamers against aggregation-prone proteins. These same aptamers can be employed as a highly specific and sensitive tool for identifying pathological FUS, with added value in revealing nuclear features invisible to antibodies. The ability of the aptamers to discriminate between physiological and pathological forms of FUS could lead to better understanding of disease mechanisms, particularly regarding nuclear and cytoplasmic aggregation dynamics.

These findings lay the groundwork for broader application in neurodegenerative diagnostics and for expanding aptamer use to other protein aggregation disorders. Nevertheless, several limitations of this study must be acknowledged. The aptamers were designed against the RNA-binding region of FUS, and although aggregation is expected to partially preserve these domains, it remains possible that certain aggregate conformations or post-translational modifications may reduce binding efficiency. Additionally, while we demonstrated specificity in post-mortem brain tissue, the behavior of these aptamers in more complex biological environments, such as living tissue or biofluids, remains to be fully assessed. Moreover, although specificity towards FUS was prioritized computationally, cross-reactivity with other RNA-binding proteins sharing similar domains cannot be entirely excluded and warrants further investigation. Finally, the histological application of aptamers, while powerful, will require optimization and standardization before it can be integrated into routine diagnostic workflows.

## MATERIALS and METHODS

Aptamer design. We targeted the RNA-binding region of FUS (residues 269–454: RRM, RGG2, ZnF) to exploit its natural nucleic-acid affinity. High-confidence FUS-binding sites were called from ENCODE eCLIP data (ENCODE project ^55^) (HepG2 & K562; two replicates each) after filtering for log₂(fold-enrichment) > 3 (Fig. 1A). Merging all four replicates yielded 4,203 reproducible binding sites; To measure the ability of catRAPID to reproduce the experimental data, we used these eCLIP binding sites as the positive set and an equally sized random selection of sites from the negative set as controls. A receiver operating characteristic (ROC) curve was computed to assess the ability of catRAPID to distinguish true FUS-binding sites from non-binding regions. The resulting area under the curve (AUC) was 0.868, indicating strong predictive performance (Fig. 1B).

Of the eCLIP sites, 401 of these overlapped with PAR-CLIP ^56^ and were designated as our positive set for the motif detection (Fig. 1A). As controls, we pooled all eCLIP peaks seen in only one replicate, failing the same enrichment cutoff and non-overlapping with PAR-CLIP, giving 200,664 sites. From these we randomly sampled 1,075 sequences to create a negative input set that was matching the length distribution of the positive set. Both sets were introduced to STREME (MEME Suite) for de novo motif discovery (motif lengths 4–16 nt), identifying eight enriched motifs (5–13 nt) in the positive set. These were supplemented by 3 FUS-recognized motifs from the catRAPID omics v2 library ^25^, generating a final set of 11 motifs. 28,102 candidate aptamers were generated by extracting RNA sequences centered around these motif occurrences, with an additional 0–20 nucleotide 31 extension. Secondary structure predictions were performed using RNAfold ^28^, and a custom scoring function was developed to prioritize aptamers displaying a perfect hairpin structure at the 51 end followed by an unstructured 31 extension. In parallel, the sequences were evaluated for the presence of at least one GGU motif in the single-stranded tail, a known enhancer of binding to FUS’ Zn finger domain ^56–58^ (Fig. 1C-D)

Sequences were further prioritized based on their predicted binding propensity to FUS_269–454_ using catRAPID ^17,25^. Aptamers with a secondary structure score above 0.9 and favorable catRAPID RNA Fitness scores were retained for experimental validation. This yielded 83 high-confidence aptamers fulfilling all criteria, while low-scoring sequences were retained as negative controls. To complement the de novo approach, several aptamers were directly designed based on high catRAPID binding predictions, favoring structured hairpins with enriched sequence motifs, even when they were not directly extracted from the CLIP datasets.

Protein purification. Purification of RRM12-RGG-ZnfC (FUS_269-454_) was carried out from kanamycin-resistant pET-Sumo expression vectors with a six-histidine tag (his-tag) and expressed in Rosetta (DE3) E. coli cells. Cells transformed with the plasmid were grown overnight at 37°C in LB medium containing 50 μg/ml kanamycin. Cell cultures were diluted 1:100 in fresh LB with 50 μg/ml kanamycin and grown until an optical density of ca. 0.6, before adding 0.5 mM IPTG and 0,1 mM ZnCl_2_ to induce protein expression overnight at 18°C. Cells were collected by centrifugation at 4800 g for 20 min at 4°C, then resuspended in lysis buffer (20 mM potassium phosphate buffer pH 7.2, 150 mM KCl, 5 mM imidazol, 0,1 mM ZnCl_2_, 1 mM DTT, 5 % v/v glycerol, 1 mg/ml lysozyme, cOmplete™ EDTA-free Protease Inhibitor, 1 μg/ml DNase I). Cells were broken by probe-sonication and soluble proteins were recovered by centrifugation at 18.000 g for 1 h at 4°C. The total soluble protein mixture was loaded onto a column packed with nickel-coated resin and the 6xHis-SUMO-construct was eluted with high-salt phosphate buffer (20 mM potassium phosphate buffer pH 7.2, 150 mM KCl, 5 uM ZnCl_2_, 1 mM DTT, 5 % v/v glycerol) with the addition of 300 mM imidazole. The eluate was dialyzed overnight at 4°C against low-salt phosphate buffer (20 mM potassium phosphate pH 7.2, 15 mM KCl, 5 uM ZnCl_2_, 1 mM DTT, 5 % v/v glycerol) in the presence of the Tobacco Etch Virus (TEV) protease at a 1:20 protein construct:TEV molar ratio, to remove the 6×His-SUMO tag. Following a further affinity purification step with a nickel-coated resin, a HiTrap Heparin column was used to remove nucleic acids. The protein was then eluted with 1.5 M KCl and submitted to size-exclusion chromatography with a HiLoad 16/60 Superdex 75 column. Protein identity and purity were assessed by PAGE. The proteins were flash frozen and stored in low salt buffer (20 mM potassium phosphate pH 7.2, 15 mM KCl, 5 uM ZnCl_2_, 1 mM DTT) at −80 °C.

Quantification of the aptamer binding constants. Biolayer interferometry experiments were acquired on an Octet Red instrument (ForteBio, Inc., Menlo Park, CA) operating at 25 °C. The binding assays were performed in 20 mM potassium phosphate buffer (pH 7.2) with 150 mM KCl and 0.05% Tween20. Streptavidin-coated biosensors were loaded with 1-5 µg/ml RNA aptamer modified with biotin on one of the ends and exposed to increasing protein concentrations varying from 101nM to 101µM, according to the strength of the binding. K_d_ values were estimated by fitting the response intensity (shift in the wavelength upon binding) as a function of the protein concentration, at the steady state. The assay was repeated at least 3 times, each time in triplicate.

Circular dichroism measurements. Circular dichroism spectra were recorded on a JASCO-1100 spectropolarimeter. The spectra were acquired in 0.1 mm path length quartz cuvettes under a constant N_2_ flush at 4.0 L/min. All CD datasets were an average of ten scans. For the determination of the global secondary structure, the far-UV (200–320 nm) spectrum was recorded at 30°C in 20 mM potassium phosphate buffer at pH 7.2 with 150 mM KF. Samples were also gradually heated to 70°C (1°C/min) for the determination of their structural stability. The ellipticity variation as expressed by the intensity of the band at 222 nm was followed as a function of temperature. The spectra were corrected for the buffer signal and expressed as mean residue molar ellipticity θ (deg1cm^2^1dmol^−1^).

Computational model predictions: RNA aptamers and protein. The RNA aptamers models were predicted by means of RNAComposer ^33,34^. This fully automated RNA prediction modeling server is able to use a fragment database dictionary, namely RNA FRABASE ^59^, to put the different RNA secondary structure elements of the RNA molecule of interest in relation with experimentally resolved tertiary structures, thus producing a 3-dimensional model. To build the prediction of aggregated FUS, we took advantage of the AlphaFold 3 AI model accuracy in predicting biomolecular interactions ^60^; using the AlphaFold Server (https://alphafoldserver.com/about) it is possible to combine multiple copies of the full-length protein (8 FUS copies, considering the web server upper limit) in the presence of an equivalent number of Zn^2+^ ions. Three independent jobs, each producing five different predictions, were run on the platform using a different randomly generated seed to increase diversity.

Protein aggregation assay. Purified FUS construct (FUS_269-454_) stored at −80°C was thawed on ice prior to the assay, and the contents were transferred to a fresh tube to ensure removal of degraded protein. The salt concentration was adjusted to a final concentration of 150 mM (from 15 mM) KCl using the storage buffer modified to contain a high concentration of salt (3 M KCl in 20 mM potassium phosphate pH 7.2, 5 μM ZnCl2, and 1 mM DTT). The protein concentration was determined by measuring absorbance at 280 nm (extinction coefficient, 1 = 20970 M^−1^1cm^−1^), and the solution adjusted to a final protein concentration of 15 µM by diluting in the aggregation buffer (20 mM potassium phosphate pH 7.2, 150 mM KCl, 5 μM ZnCl2, 1 mM DTT, and 0.01% sodium azide). The aggregation was carried out at 37°C in a 96-well plate and imaged in a plate reader (BioTek Cytation 3), with a single borosilicate glass bead in each well with the sample. The aggregation was performed with 5 replicates or wells, either with or without ProteoStat (Enzo Life Sciences), as its spectral range overlaps with that of Atto590 on the aptamers. ProteoStat was used at a final dilution of 1:1000 as per the manufacturer’s instructions. Parameters were set so that the plate reader took a fluorescence reading for each well (excitation 550 nm, emission 600 nm) every 15 minutes, at a gain of 50, with continuous double orbital shaking at 205 cpm. Aliquots were taken out at specific time points (0, 2.5, 5, 10, and 24 hours), and imaged, with either one of the aptamers or ProteoStat.

TIRF microscopy. Single-molecule imaging was carried out on an ONI Nanoimager (Oxford Nanoimaging Ltd., Oxford) equipped with a 100×/1.4 numerical aperture oil immersion apochromatic objective lens and ORCA-Flash 4.0 V3 scientific complementary metal-oxide semiconductor camera. Samples incubated with the Atto590-tagged aptamers were exposed to 561 nm excitation by total internal reflection using a laser illumination angle of 53.5°. Each field of view (FOV) was imaged for 2,000 frames at a rate of 20 frames/s, and 3 x 500 μm–spaced FOVs were captured for each well, aptamer, and timepoint to account for any region-specific variations.

Preparation of samples and aptamers for SR imaging. 24 x 60 mm borosilicate glass coverslips were cleaned by exposure to argon plasma for 45 minutes. Self-adhesive 18-well Ibidi chamber slides were affixed onto plasma-cleaned coverslips. FUS aggregates were diluted 1:4 in the aggregation buffer and allowed to adsorb onto coverslips for 1 hour, then washed three times with PBS buffer (0.02 μm-filtered, Anotop25, Whatman). Atto590-tagged aptamers were diluted in filtered PBS and used at a final concentration of 5 nM for imaging.

SR image analysis. The positions of the transiently immobilized aptamers within each frame were determined using the ImageJ/Fiji plugin, PeakFit, of the GDSC Single Molecule Light Microscopy package (http://www.sussex.ac.uk/gdsc/intranet/microscopy/imagej/gdsc_plugins). A signal strength threshold of 30 and a precision threshold of 20 was used. The localizations were grouped into clusters using the DBSCAN algorithm in Python 3.8 (sklearn v0.24.2) using epsilon=10.5 pixels and a minimum localisations threshold of 50 to remove random localisations. Clustered localisations were plotted as 2D Gaussian distributions, with a width equal to the precision that they were localised to. For the length of each cluster, the localisations were plotted with widths equal to the precision FWHM and were then analysed using the measure module (skimage v0.18.1). The lengths plotted are the major axis length.

Post-fixation Cellular Staining with Aptamers. Human neuroblastoma (SK-N-BE) cells were cultured in RPMI medium, and maintained at 37 °C in a humidified incubator with 5% CO₂. For microscopy studies, cells were seeded in 24-well plates containing coverslips pre-treated with an attaching factor (ThermoFisher Scientific, S006100). After 24 h, when confluence reached approximately 65%, protein expression was induced by addition of 100 ng/ml doxycycline for either the wild-type (WT) or P525L mutant protein. After an additional 24 h, cells were stressed with 0.5 mM sodium arsenite (SA) for 1 h at 37 °C. Following treatment, cells were fixed with 4% paraformaldehyde (PFA) for subsequent staining procedures. Permeabilization was performed using either 5 µM digitonin in PBS for 5 min or 0.5% Triton X-100 in PBS for 10 min at room temperature (rT). To reduce nonspecific binding, cells were blocked for 1 h at rT in PBS containing 5% bovine serum albumin (BSA) and 100 µg/mL total yeast RNA. For antibody staining, cells were incubated with a primary antibody, anti-FUS/TSL (Santa Cruz, sc-47711, 1:250), diluted in blocking solution (5% BSA in PBS) for 1 h at rT, followed by three washes with PBS. The secondary antibody, goat anti-mouse Alexa Fluor 488 (ThermoFisher Scientific, A11029, 1:500), diluted in the same blocking solution was then applied for 1 h at rT and removed by three subsequent PBS washes. Aptamer staining was performed by incubating cells with 150 nM fluorophore-conjugated aptamer (Atto 590) in a blocking solution containing 100 µg/mL total yeast RNA in PBS (without BSA) for 1 h at rT. Nuclei were counterstained with 0.5 µg/mL 41,6-diamidino-2-phenylindole (DAPI) solution for 5 min, followed by three final washes in PBS. Coverslips were mounted onto glass slides using a drop of mounting medium ProLong™ Diamond Antifade Mountant (Invitrogen).

Images acquisition and analysis. Glass slides with fixed cells were visualized with a Nikon’s A1R MP multiphoton confocal microscope, employing the ×60 objective and 3 channel non-descanned detectors. Image analyses were performed with a custom Python pipeline implemented using tifffile, scikit-image, OpenCV, and NumPy for quantitative analysis of multichannel TIFF microscopy images. Images were loaded and user-defined channel indices were assigned for nuclei, cytoplasmic/cell signal, and one or two condensate channels. Nuclear segmentation was performed by Gaussian smoothing followed by Otsu thresholding, morphological closing, hole filling, and removal of small objects. Cell boundaries were reconstructed via a marker-controlled watershed, using nuclear centroids as markers and a blurred cytoplasmic channel to define the segmentation mask. Condensate detection was performed by subtracting a strong Gaussian background estimate (σ=20) from the raw aggregate channel, thresholding relative to the denoised maximum intensity, and filtering by object area and normalized mean intensity. Condensates touching nuclei were excluded. Each remaining condensate was assigned to the cell with the highest pixel-level overlap, and colocalization between condensate channels was computed as the spatial intersection of their respective masks after nuclear exclusion. For each cell, the script exported a CSV file containing condensate counts, colocalized events, and per-cell condensate mean intensities. Additionally, RGB overlays were generated to visualize nuclear and cellular boundaries together with condensate detections. A batch-level summary table reporting total condensate, colocalized condensate, and global condensate intensity metrics was also produced.

Cytosolic and nuclear protein extraction. Cell pellets were resuspended in 400 µL cell lysis buffer (5 mM PIPES pH 8.0, 85 mM KCl, 0.5% NP-40) per 10 million cells and incubated on ice for 10 min. Cells were homogenized on ice using a dounce homogenizer with a tight pestle by applying 20 strokes, pausing for 2 min, and repeating with an additional 20 strokes. The homogenate was transferred to a new 1.5 mL tube, and nuclei were pelleted by centrifugation at 1500 g for 5 min at 4 °C. The supernatant was transferred in a new tube as this constitutes the cytosolic fraction. Nuclei were resuspended in 100 µL nuclear lysis buffer (25 mM Tris pH 8.1, 5 mM EDTA, 0.5% SDS) per 10 million cells and incubated on ice for 10 min. The suspension was aliquoted into 100 µL portions in sonication tubes and processed using a Bioruptor Pico sonicator for 5–7 cycles of 30 s on/30 s off, ensuring the device was pre-cooled for at least 20 min (to 7–8 °C). Nuclear extract was flash-frozen and stored at −20 °C.

The purity of cytosolic and nuclear fractions was assessed by western blot. Membranes were probed with the following primary antibodies: anti-FUS/TSL (Santa Cruz, sc-47711, 1:500), anti-vinculin (Abcam, ab91459, 1:1000) as a cytosolic housekeeping control, and anti-histone H3 (Active Motif, 61647, 1:1000) as a nuclear housekeeping control. Membranes were then incubated with species-appropriate HRP-conjugated secondary antibodies: goat anti-rabbit IgG-HRP (ThermoFisher Scientific, 31460, 1:10000) or goat anti-mouse IgG-HRP (BioRad, 1706516, 1:10000). The peroxidase substrate was added (Clarity Western ECL Substrate) and signal detection was performed using a ChemiDoc imaging system (Bio-Rad).

Human post-mortem tissue staining. All postmortem data were generated from tissue samples collected and banked at the New York Brain Bank of Columbia University with consent obtained from the patient’s next of kin, according to New York State law and the guidelines of the Department of Pathology of Columbia University and New York Presbyterian Hospital. In the State of New York, research involving autopsy material does not meet the regulatory definition of ‘human subject research’ and is not subject to institutional review board oversight. Formalin-fixed paraffin embedded (FFPE) tissue from BA4 was sectioned on a Leica microtome in 4 µm sections onto a superfrost microscope slide. Sections were dried overnight at 40 °C and immunostaining was performed using the Novolink Polymer detection system with an anti-FUS antibody (BioTechne NB100-595) at a 1 in 500 dilution. DAB chromogen was used and counterstaining was performed with haematoxylin, according to standard operating procedures. Pretreatment was 30 minutes in a pressure cooker in pH6 buffered citric acid. FUS aptamer staining was performed as previously published using identical method and concentration as TDP-43 aptamer staining ^61^. Digital burden scoring was performed using the freely available QuPath software implementing superpixel analysis using the following code: setImageType(’BRIGHTFIELD_H_DAB’); setColorDeconvolutionStains(’{“Name” : “H-DAB default”, “Stain 1” : “Hematoxylin”, “Values 1” : “0.65111 0.70119 0.29049 ”, “Stain 2” : “DAB”, “Values 2” : “0.26917 0.56824 0.77759 ”, “Background” : ” 255 255 255 “}’); setPixelSizeMicrons(0.625,0.625); createSelectAllObject(true); selectAnnotations (); runPlugin(’qupath.imagej.superpixels.DoGSuperpixelsPlugin’, ’{“downsampleFactor”: 1.0, “sigmaMicrons”: 3, “minThreshold”: 10.0, “maxThreshold”: 230.0, “noiseThreshold”: 0.0}’); selectDetections(); runPlugin(’qupath.lib.algorithms.IntensityFeaturesPlugin’, ’{“downsample”: 1.0, “region”: “ROI”, “tileSizePixels”: 200.0, “colorOD”: false, “colorStain1”: false, “colorStain2”: true, “colorStain3”: false, “colorRed”: false, “colorGreen”: false, “colorBlue”: false, “colorHue”: false, “colorSaturation”: false, “colorBrightness”: false, “doMean”: true, “doStdDev”: false, “doMinMax”: false, “doMedian”: false, “doHaralick”: false, “haralickDistance”: 1, “haralickBins”: 32}’); setDetectionIntensityClassifications(“ROI: 2.00 µm per pixel: DAB: Mean”, 0.2689, 0.3007, 1). H-Score was calculated using the following weighted score (1x number of 1+ detections + 2x number of 2+ detections + 3x number of 3+ detections) normalised by the total number of pixels in the region of interest. Graphs were plotted using GraphPad Prism v10.0. Mann Whitney U test (with correction for multiple testing) was used to compare between cases and stains.

## Acknowledgements

We thank the “RNA Flagship” at IIT, as well as all members of the J.M.G. and G.G.T. research groups, for their support and contributions. A special thanks to Dr. Alessio Colantoni for the help with the analysis of the CLIP data. The research leading to this work was supported by the ERC ASTRA_855923 (G.G.T.), EIC Pathfinder IVBM4PAP_101098989 (G.G.T.) and PNRR grant from the National Centre for Gene Therapy and Drugs based on RNA Technology (CN00000041 EPNRRCN3 (G.G.T.). F.D.P. acknowledges the support granted by the European Union—Next Generation EU, Mission 4 Component 1 CUP D53D23016360001, PRIN-PNRR, Grant n. P2022CLXMK, for his current position. The work was also funded by a grant from Target ALS (BB-2022-C4-L2) to MHH, NS, GGT, EZ, and JMG employing FMW, ADS, and MG. Also an MND Primer award from LifeArc (11.12.2023) to JMG and MHH employing RJ.

## References

1. Kodavati, M. et al. FUS unveiled in mitochondrial DNA repair and targeted ligase-1 expression rescues repair-defects in FUS-linked motor neuron disease. Nat. Commun. 15, 2156 (2024).

2. Errichelli, L. et al. FUS affects circular RNA expression in murine embryonic stem cell-derived motor neurons. Nat. Commun. 8, 14741 (2017).

3. Reber, S. et al. Minor intron splicing is regulated by FUS and affected by ALS-associated FUS mutants. EMBO J. 35, 1504–1521 (2016).

4. Naumann, M. et al. Phenotypes and malignancy risk of different FUS mutations in genetic amyotrophic lateral sclerosis. Ann. Clin. Transl. Neurol. 6, 2384–2394 (2019).

5. López-Erauskin, J. et al. ALS/FTD-linked mutation in FUS suppresses intra-axonal protein synthesis and drives disease without nuclear loss-of-function of FUS. Neuron 100, 816–830.e7 (2018).

6. Nolan, M., Talbot, K. & Ansorge, O. Pathogenesis of FUS-associated ALS and FTD: insights from rodent models. Acta Neuropathol. Commun. 4, 99 (2016).

7. Naumann, M. et al. Impaired DNA damage response signaling by FUS-NLS mutations leads to neurodegeneration and FUS aggregate formation. Nat. Commun. 9, 335 (2018).

8. Scekic-Zahirovic, J. et al. Cytoplasmic FUS triggers early behavioral alterations linked to cortical neuronal hyperactivity and inhibitory synaptic defects. Nat. Commun. 12, 3028 (2021).

9. Mackenzie, I. R. A. & Neumann, M. Fused in Sarcoma Neuropathology in Neurodegenerative Disease. Cold Spring Harb. Perspect. Med. 7, (2017).

10. Rezvykh, A. P. et al. Cytoplasmic aggregation of mutant FUS causes multistep RNA splicing perturbations in the course of motor neuron pathology. Nucleic Acids Res. 51, 5810–5830 (2023).

11. Nomura, T. et al. Intranuclear aggregation of mutant FUS/TLS as a molecular pathomechanism of amyotrophic lateral sclerosis. J. Biol. Chem. 289, 1192–1202 (2014).

12. Tacconelli, S., et al. A validated panel of commercial antibodies for reliable detection of FET proteins. bioRxiv (2025) doi:10.1101/2025.11.19.689317.

13. Alshalfie, W. et al. The identification of high-performing antibodies for RNA-binding protein FUS for use in Western Blot, immunoprecipitation, and immunofluorescence. F1000Res. 12, 376 (2023).

14. Shneider, N. A. et al. Antisense oligonucleotide jacifusen for FUS-ALS: an investigator-initiated, multicentre, open-label case series. Lancet 405, 2075–2086 (2025).

15. Cox, D. et al. Quantitative profiling of nanoscopic protein aggregates reveals specific fingerprint of TDP-43-positive assemblies in motor neuron disease. Adv. Sci. (Weinh.) 12, e05484 (2025).

16. Ni, S. et al. Recent progress in aptamer discoveries and modifications for therapeutic applications. ACS Appl. Mater. Interfaces 13, 9500–9519 (2021).

17. Zacco, E. et al. Probing TDP-43 condensation using an in silico designed aptamer. Nat. Commun. 13, 3306 (2022).

18. Ellington, A. D. & Szostak, J. W. In vitro selection of RNA molecules that bind specific ligands. Nature 346, 818–822 (1990).

19. Lerga, A. et al. Identification of an RNA binding specificity for the potential splicing factor TLS. J. Biol. Chem. 276, 6807–6816 (2001).

20. Wang, X., Schwartz, J. C. & Cech, T. R. Nucleic acid-binding specificity of human FUS protein. Nucleic Acids Res. 43, 7535–7543 (2015).

21. Imperatore, J. A., McAninch, D. S., Valdez-Sinon, A. N., Bassell, G. J. & Mihailescu, M. R. FUS Recognizes G Quadruplex Structures Within Neuronal mRNAs. Front Mol Biosci 7, 6 (2020).

22. Loughlin, F. E. et al. The Solution Structure of FUS Bound to RNA Reveals a Bipartite Mode of RNA Recognition with Both Sequence and Shape Specificity. Mol. Cell 73, 490–504.e6 (2019).

23. Milordini, G. et al. Computationally-designed aptamers targeting RAD51-BRCA2 interaction impair homologous recombination and induce synthetic lethality. Nat. Commun. 1–18 (2025).

24. Luige, J. et al. Design and characterization of G-quadruplex RNA aptamers reveal RNA-binding by KDM5 lysine demethylases. Comput. Struct. Biotechnol. J. 27, 2719–2729 (2025).

25. Armaos, A., Colantoni, A., Proietti, G., Rupert, J. & Tartaglia, G. G. catRAPID omics v2.0: going deeper and wider in the prediction of protein-RNA interactions. Nucleic Acids Res. 49, W72–W79 (2021).

26. Rupert, J., Monti, M., Zacco, E. & Tartaglia, G. G. RNA sequestration driven by amyloid formation: the alpha synuclein case. Nucleic Acids Res. 51, 11466–11478 (2023).

27. Jutzi, D. et al. Aberrant interaction of FUS with the U1 snRNA provides a molecular mechanism of FUS induced amyotrophic lateral sclerosis. Nat. Commun. 11, 6341 (2020).

28. Lorenz, R. et al. ViennaRNA Package 2.0. Algorithms Mol. Biol. 6, 26 (2011).

29. Loughlin, F. E. & Allain, F. H.-T. Solution structure of FUS-RRM bound to stem-loop RNA. Preprint at 10.2210/pdb6gbm/pdb (2019).

30. Loughlin, F. E. & Allain, F. H.-T. Solution structure of FUS-ZnF bound to UGGUG. Preprint at 10.2210/pdb6g99/pdb (2019).

31. Schwartz, J. C. et al. FUS binds the CTD of RNA polymerase II and regulates its phosphorylation at Ser2. Genes Dev. 26, 2690–2695 (2012).

32. Luige, J., Armaos, A., Tartaglia, G. G. & Ørom, U. A. V. Predicting nuclear G-quadruplex RNA-binding proteins with roles in transcription and phase separation. Nat. Commun. 15, 2585 (2024).

33. Popenda, M. et al. Automated 3D structure composition for large RNAs. Nucleic Acids Res. 40, e112 (2012).

34. Sarzynska, J., Popenda, M., Antczak, M. & Szachniuk, M. RNA tertiary structure prediction using RNAComposer in CASP15. Proteins 91, 1790–1799 (2023).

35. Kypr, J., Kejnovská, I., Renciuk, D. & Vorlícková, M. Circular dichroism and conformational polymorphism of DNA. Nucleic Acids Res. 37, 1713–1725 (2009).

36. Horrocks, M. H. et al. Single-molecule imaging of individual amyloid protein aggregates in human biofluids. ACS Chem. Neurosci. 7, 399–406 (2016).

37. Saleeb, R. S. et al. Two-color coincidence single-molecule pulldown for the specific detection of disease-associated protein aggregates. Sci. Adv. 9, eadi7359 (2023).

38. Banterle, N., Bui, K. H., Lemke, E. A. & Beck, M. Fourier ring correlation as a resolution criterion for super-resolution microscopy. J. Struct. Biol. 183, 363–367 (2013).

39. Lo Bello, M., et al. ALS-related mutant FUS protein is mislocalized to cytoplasm and is recruited into stress granules of fibroblasts from asymptomatic FUS P525L mutation carriers. Neurodegener. Dis. 17, 292–303 (2017).

40. Mariani, D. et al. ALS-associated FUS mutation reshapes the RNA and protein composition of stress granules. Nucleic Acids Res. 52, 13269–13289 (2024).

41. Adam, S. A., Sterne-Marr, R. & Gerace, L. Nuclear protein import using digitonin-permeabilized cells. Methods Enzymol. 219, 97–110 (1992).

42. Marrone, L. et al. Isogenic FUS-eGFP iPSC reporter lines enable quantification of FUS stress granule pathology that is rescued by drugs inducing autophagy. Stem Cell Reports 10, 375–389 (2018).

43. Baron, D. M. et al. Amyotrophic lateral sclerosis-linked FUS/TLS alters stress granule assembly and dynamics. Mol. Neurodegener. 8, 30 (2013).

44. Korobeynikov, V. A., Lyashchenko, A. K., Blanco-Redondo, B., Jafar-Nejad, P. & Shneider, N. A. Antisense oligonucleotide silencing of FUS expression as a therapeutic approach in amyotrophic lateral sclerosis. Nat. Med. 28, 104–116 (2022).

45. Spence, H. et al. RNA aptamer reveals nuclear TDP-43 pathology is an early aggregation event that coincides with STMN-2 cryptic splicing and precedes clinical manifestation in ALS. Acta Neuropathol. 147, 50 (2024).

46. Bankhead, P. et al. QuPath: Open source software for digital pathology image analysis. Sci. Rep. 7, 16878 (2017).

47. Zacco, E. et al. RNA as a key factor in driving or preventing self-assembly of the TAR DNA-binding protein 43. J. Mol. Biol. 431, 1671–1688 (2019).

48. Shelkovnikova, T. A., Robinson, H. K., Southcombe, J. A., Ninkina, N. & Buchman, V. L. Multistep process of FUS aggregation in the cell cytoplasm involves RNA-dependent and RNA-independent mechanisms. Hum. Mol. Genet. 23, 5211–5226 (2014).

49. Germer, K., Leonard, M. & Zhang, X. RNA aptamers and their therapeutic and diagnostic applications. Int. J. Biochem. Mol. Biol. 4, 27–40 (2013).

50. Horrocks, M. H., Palayret, M., Klenerman, D. & Lee, S. F. The changing point-spread function: single-molecule-based super-resolution imaging. Histochem. Cell Biol. 141, 577–585 (2014).

51. Whiten, D. R. et al. Nanoscopic characterisation of individual endogenous protein aggregates in human neuronal cells. Chembiochem 19, 2033–2038 (2018).

52. Shi, S.-R., Shi, Y. & Taylor, C. R. Antigen retrieval immunohistochemistry: review and future prospects in research and diagnosis over two decades. J. Histochem. Cytochem. 59, 13–32 (2011).

53. Grassano, M. et al. Phenotype analysis of fused in sarcoma mutations in amyotrophic lateral sclerosis. Neurol. Genet. 8, e200011 (2022).

54. Aladesuyi Arogundade, O., et al. Nucleolar stress in C9orf72 and sporadic ALS spinal motor neurons precedes TDP-43 mislocalization. Acta Neuropathol. Commun. 9, 26 (2021).

55. Van Nostrand, E. L. et al. A large-scale binding and functional map of human RNA-binding proteins. Nature 583, 711–719 (2020).

56. De Santis, R. et al. Mutant FUS and ELAVL4 (HuD) Aberrant Crosstalk in Amyotrophic Lateral Sclerosis. Cell Rep. 27, 3818–3831.e5 (2019).

57. Bellucci, M., Agostini, F., Masin, M. & Tartaglia, G. G. Predicting protein associations with long noncoding RNAs. Nat. Methods 8, 444–445 (2011).

58. Dominguez, D. et al. Sequence, structure, and context preferences of human RNA binding proteins. Mol. Cell 70, 854–867.e9 (2018).

59. Popenda, M. et al. RNA FRABASE 2.0: an advanced web-accessible database with the capacity to search the three-dimensional fragments within RNA structures. BMC Bioinformatics 11, 231 (2010).

60. Abramson, J. et al. Accurate structure prediction of biomolecular interactions with AlphaFold 3. Nature 630, 493–500 (2024).

61. M Waldron, F., Spence, H. & Gregory, J. TDP-43 RNA aptamer staining to detect pathological TDP-43 in FFPE human tissue, as described in Spence and Waldron et al., 2024 (Acta Neuropathologica): A SOP and tick-sheet. v2. v2. (2024) doi:10.17504/protocols.io.eq2lyjo4mlx9/v2.

