## Supplementary Material for "Computationally Designed RNA Aptamers Enable Selective Detection of FUS Pathology in ALS"

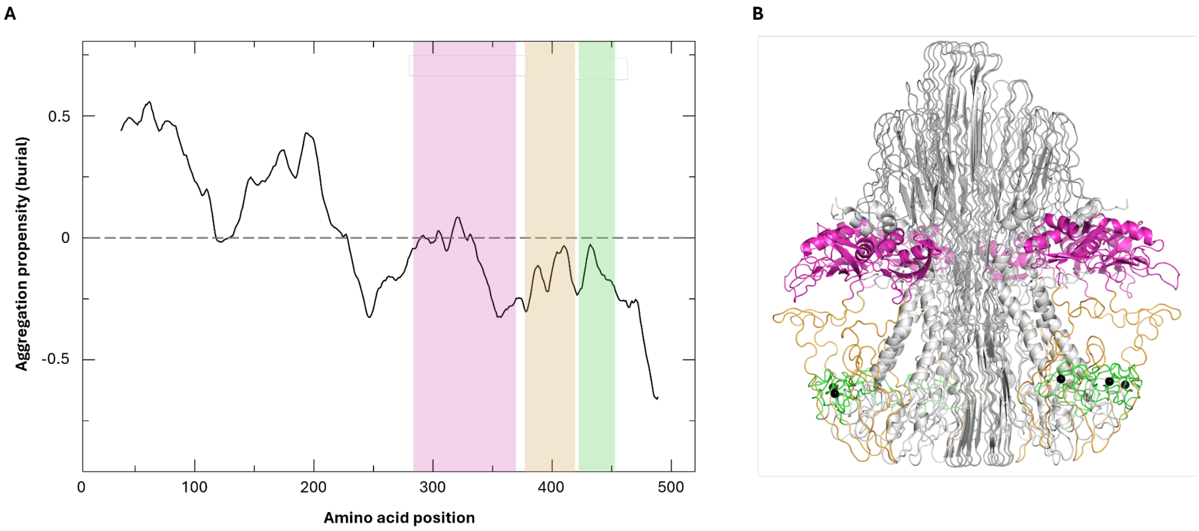


**Suppl. Fig. 1: The RNA-binding regions of FUS are predicted to be external to the aggregation core and thus available for aptamer binding. A**: Zyggregator calculation [1,2] of the residues most prone to aggregate and, therefore, to be buried in the core. The protein region starting from amino acid 210 to the C-terminus is predicted to be exposed to the solvent and ready for interaction. **B**: AlphaFold reconstruction [3] of an aggregate formed by eight copies of FUS, showing that indeed the RRM (magenta), RGG2 (yellow) and zinc finger (green) are exposed to the surface of the aggregate and available to interact with RNA binding partners.

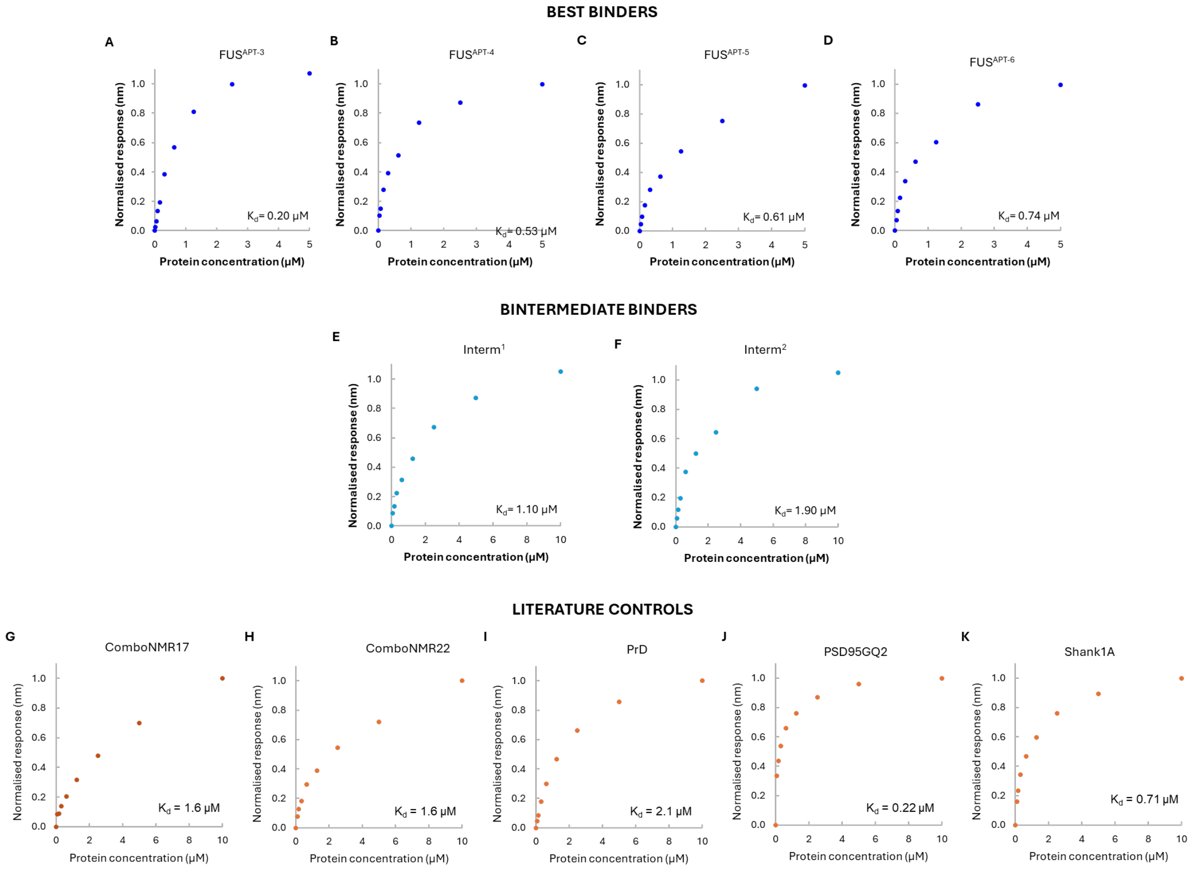


**Suppl. Fig. 2: Biolayer interferometry-derived binding curves of selected aptamers and literature controls with FUS.** Each graph reports the binding response as a function of the protein concentration and the $K_d$ values calculated at steady state.

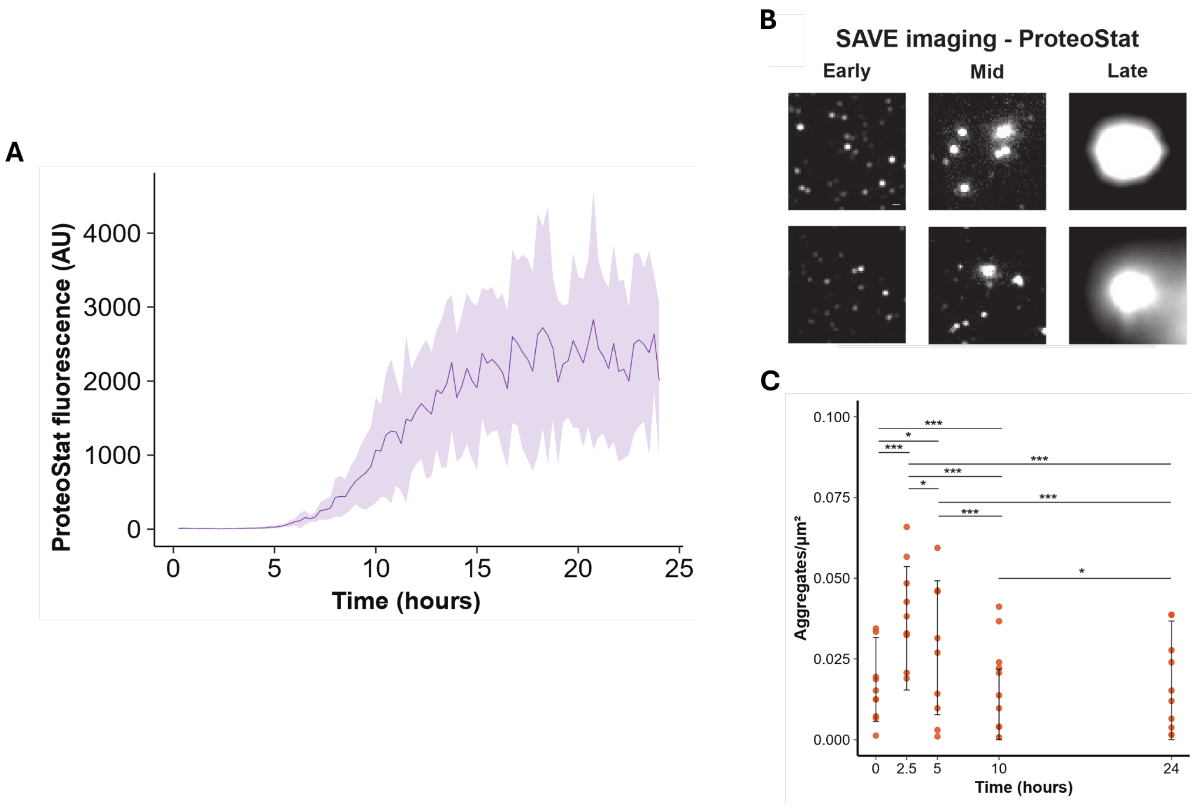


**Suppl. Fig. 3: ProteoStat fluorescence demonstrates that aggregates are formed over the 24-hour period of aggregation. A**. ProteoStat fluorescence as measured across 5 wells over time, showing an increase in average fluorescence (the solid line represents the mean, and the shaded region, the standard deviation). **B**. Representative diffraction-limited SAVE (Single-Aggregate Visualisation by Enhancement) images and **C**. quantification of the imaged aggregates formed over time. Aggregates are shown to decrease in number as they increase in size. In B, the 0 hour baseline is the "Early" timepoint, 5 hours is "Mid", and 24 hours is "Late". Scale bars = 500 nm.

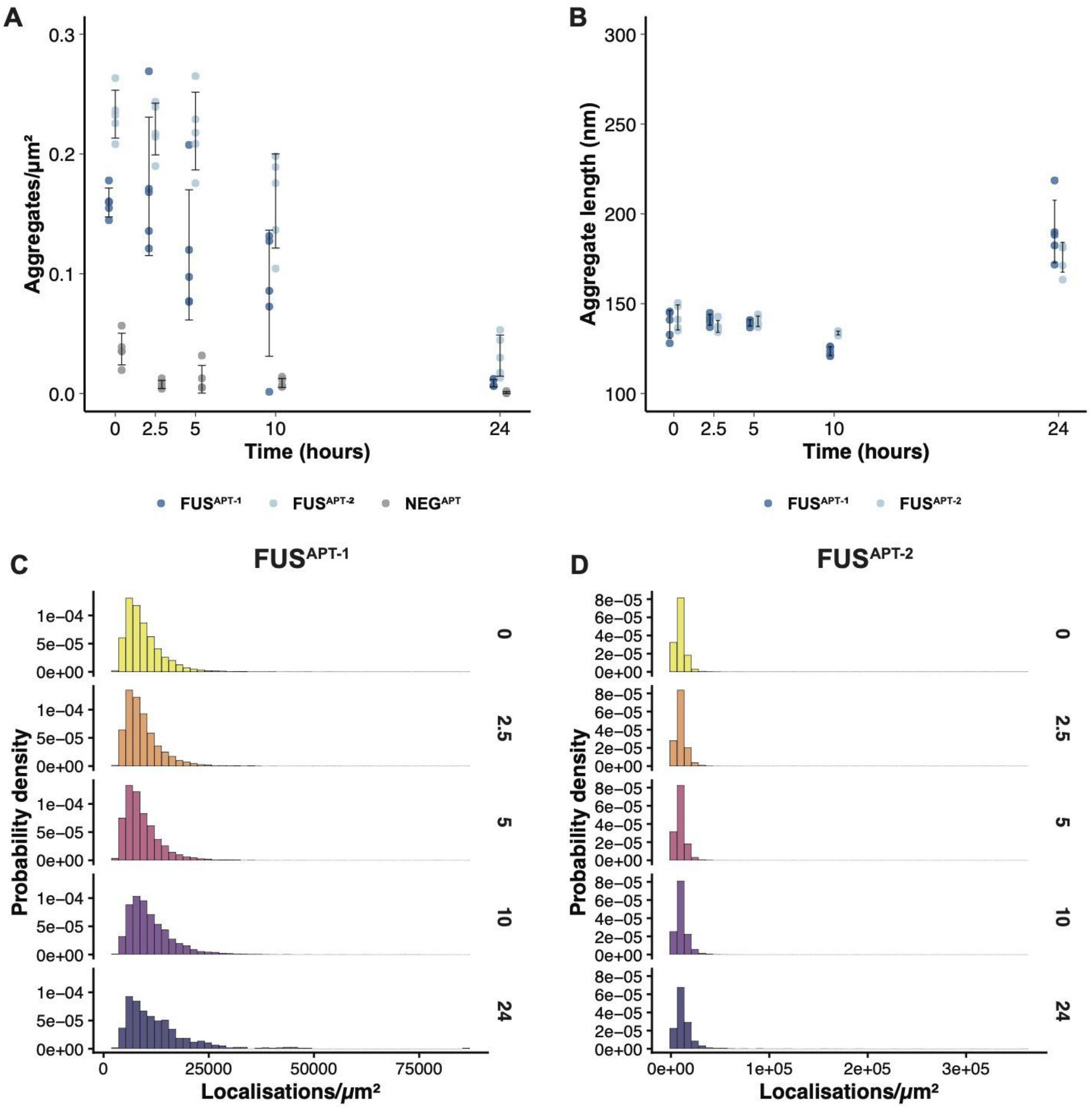


**Suppl. Fig. 4: Numbers, lengths, and localisations per unit area as detected by super-resolution imaging with the aptamers.** Quantification of **A**: numbers of aggregates detected by all 3 aptamers and **B**: lengths of the aggregates detected by the detection aptamers used in this study. Each point is the average for one well of the aggregation, for a total of 5 technical replicates per aptamer. Statistical evaluation for aggregate lengths (Kruskal-Wallis test, with Dunn post-hoc and Benjamini-Hochberg correction) resulted in multiple significant comparison, reported in **Suppl. Table 2**. Normalised histograms of the probability density distribution of localisations per unit area as detected by **C**: $FUS^{APT-1}$ or **D**: $FUS^{APT-2}$.

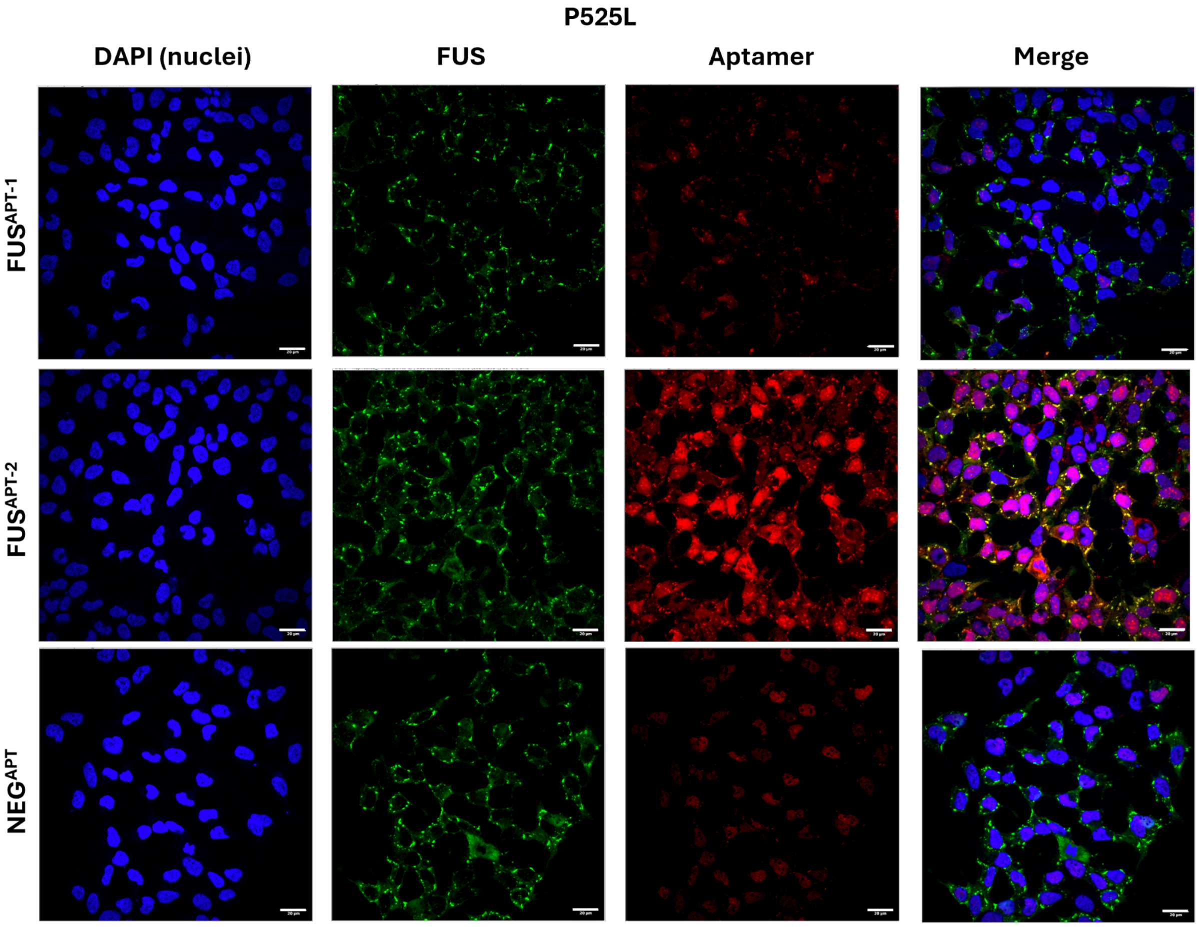


**Suppl. Fig. 5**: **Overview of immunofluorescence images of FUS P525L–expressing cells subjected to sodium arsenite stress and stained with an anti-FUS antibody together with one of the tested aptamers.** Top row: $FUS^{APT-1}$; middle row: $FUS^{APT-2}$; bottom row: $NEG^{APT}$. Nuclei are labeled with DAPI (blue); the anti-FUS primary antibody is detected using an Atto488-conjugated secondary antibody (green); aptamers are labeled with an Atto590 fluorophore (red). Scale bar= 20 µm.

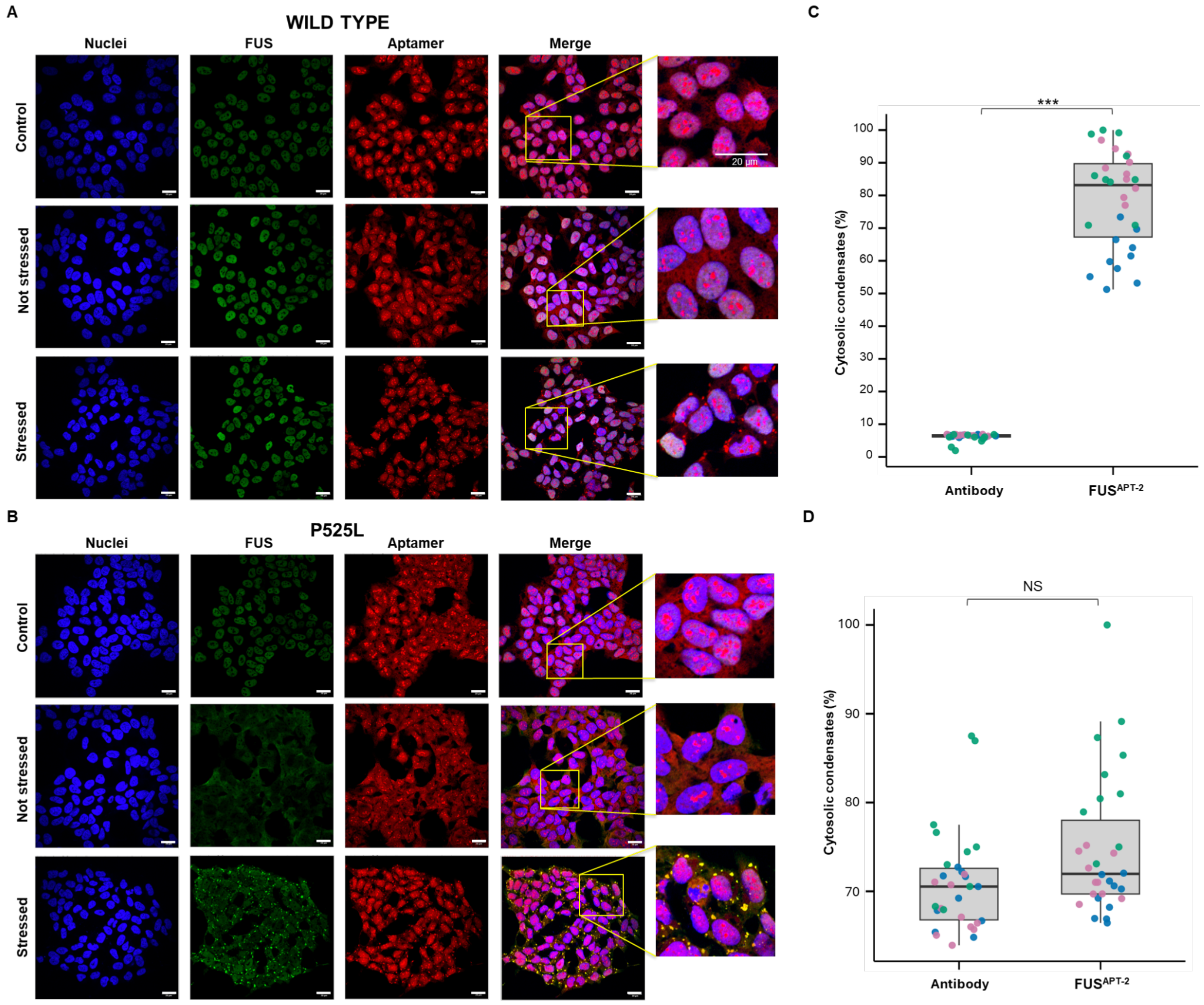


**Suppl. Fig. 6: Immunofluorescence overview of SK-N-BE cells permeabilized with Triton X-100 and stained with an anti-FUS antibody together with FUS$^{APT\text{-}2}$. A:** Cells expressing wild-type FUS. **B**: Cells expressing the FUS P525L variant. For each condition, cells are shown untreated (‘Control’, top rows), following doxycycline-induced FUS expression (‘Not stressed’, middle rows), and after exposure to sodium arsenite (‘Stressed’, bottom rows). Nuclei are labeled with DAPI (blue); the anti-FUS primary antibody is detected using an Atto488-conjugated secondary antibody (green); aptamers are labeled with an Atto590 fluorophore (red). Scale bar = 20 µm. **C–D**: Quantification of the percentage of cytosolic condensates detected by the antibody and by FUS$^{APT\text{-}2}$ in cells expressing wild-type FUS (**C**) or FUS P525L (**D**). Points represent data from triplicate experiments (blue: replicate 1; pink: replicate 2; green: replicate 3). Statistical significance was assessed using a one-tailed *t*-test: *** $p = 0.0001$.

**Suppl. Table 1**: Summary of statistical analyses for super-resolution imaging experiments (Fig. 4 and Suppl. Figs. 3–4).

### References


1. Tartaglia, G. G. *et al.* Prediction of aggregation-prone regions in structured proteins. *J. Mol. Biol.* **380**, 425–436 (2008).
2. Rupert, J., Monti, M., Zacco, E. & Tartaglia, G. G. RNA sequestration driven by amyloid formation: the alpha synuclein case. *Nucleic Acids Res.* **51**, 11466–11478 (2023).
3. Abramson, J. *et al.* Accurate structure prediction of biomolecular interactions with AlphaFold 3. *Nature* **630**, 493–500 (2024).
